# *In silico* screening of candidate NAMPT modulators for treatment of age-related diseases

**DOI:** 10.1101/2025.04.07.647561

**Authors:** Jarmila Nahálková, Fernando Berton Zanchi

## Abstract

The mitochondrial bioenergetics hypothesis postulates a critical role of mitochondrial function in aging and its deterioration is a cause of multiple age-related diseases. A superior approach to counteract age-dependent decline in cellular NAD+ levels is to activate nicotinamide phosphoribosyl transferase (NAMPT), a rate-limiting enzyme in the NAD+ salvage pathway, thereby producing the NAD^+^ precursor, nicotinamide mononucleotide. NAMPT inhibitors are also desired due to their potency in limiting cancer cell growth. Here is presented an *in-silico* screening of NAMPT modulators using compounds from traditional African, Chinese, and Russian medicinal plants and by repurposing FDA libraries. The compounds were selected by prediction to pass the blood-brain barrier, for low toxicity, and desirable pharmacokinetic and drug-likeness characteristics, and by molecular docking with crystal structures of NAMPT. A panel of 21 compounds of known NAMPT activators, inhibitors, negative controls, and randomly chosen compounds was used to validate the docking method. 2D protein-ligand interactions of the candidate compounds were critically evaluated and compared with interactions of the control panel. The virtual screening selection yielded 17 compounds with decreased predicted ΔG when bound to 6 co-crystallized structures of NAMPT. The candidates were then sorted into three binding groups based on 2D interactions with NAMPT active and allosteric sites. The stability of NAMPT complexes with stylopine, hedychilactone A, and isobavachromene, as representatives of the three binding groups, was also confirmed by molecular dynamics simulations. The results of the virtual screening should be used for experimental development of anti-aging treatments and medicines for neurodegenerative diseases and brain cancer.

## Introduction

Decades of research dedicated to mitochondrial functions led to the development of a “mitochondrial bioenergetics hypothesis,” built on a significant contribution of mitochondrial fitness to aging and age-related diseases. They contribute to the most essential cellular functions, including energy manufacture and the production and detoxification of reactive oxygen species. Mitochondria produce a proton flow by the activity of the respiratory chain complexes I, II, III, and IV located on the inner mitochondrial membrane and driven by the electron flow from NADH to oxygen, followed by the regeneration of NAD^+^. The protons exported from the mitochondrial matrix to the intermembrane space create a membrane potential, which drives their re-entry back to the mitochondrial matrix through ATP synthase, generating ATP from ADP [1]. Since the complex I of the electron transport chain (ETC) is involved in cellular energy production and NAD^+^ regeneration, it accordingly contributes to the maintenance of the NAD^+^/NADH ratio important for cellular homeostasis. The ratio decreases during the process of aging, probably due to the increased NAD^+^ consumption by PARP-s and CD38 in combination with the NAD^+^ production decline [2]. The age-dependent depletion of NAD^+^ is particularly linked to Alzheimeŕs disease (AD), Parkinsońs disease (PD), Huntingtońs disease (HD), and amyotrophic lateral sclerosis (ALS) pathologies. As evidence, the boosting of NAD^+^ levels can inhibit the occurrence of these pathologies in animal models [2]. Further, supplementing NAD^+^ precursors and/or the inhibition of NAD^+^-consuming proteins has a neuroprotective effect against the mitochondrial defects of PD models of *D. melanogaster* [3]. Increasing NAD^+^ production also positively affects 10 hallmarks of aging [2][4], which highlights the significance of NAD^+^ maintenance for healthy aging and the prevention of associated diseases.

The activation of nicotinamide phosphoribosyltransferase (NAMPT) is a superior approach to increase the cellular NAD^+^/NADH ratio and balance cellular bioenergetics decline during aging. NAMPT was originally purified and characterized in 1957 as an enzyme producing nicotinamide mononucleotide (NMN) from nicotinamide (NAM) by transfer of ribosyl pyrophosphate group of 5-phosphoribosyl 1-pyrophosphate (PPRP) [5]. NMN is then converted to NAD^+^ by nicotinamide nucleotide adenylyl transferase (NMNAT) [6][7]. NAMPT is a rate-limiting enzyme of NAD^+^ biosynthesis by salvage of NAM produced by NAD^+^ catabolism of NAD^+^-dependent proteins with NAD-ase activity, such as SIRT-s, PARP-s, and CD38 [8].

Interestingly, the anti-aging and lifespan-extending effect was achieved by supplementation of NAMPT-containing extracellular vesicles to experimental mice [9]. The relevance of the enzyme to the pathology of neurodegenerative disease was demonstrated by the deletion of NAMPT in projection neurons, which triggered mitochondrial and synaptic dysfunctions and further caused neurodegeneration specifically located in the motor cortex. It was also accompanied by muscular atrophy and gradual paralysis, loss of brown adipose tissue (BAT), and shortened lifespan, which were partially rescued by nicotinamide mononucleotide (NMN) in the diet [10]. The dopaminergic neurons are more vulnerable to NAMPT depletion, which leads to Parkinsońs disease-like symptoms in mice [11]. Homozygous NAMPT deficiency is lethal in utero; however, heterozygous animals survive relatively normally, with decreased intracellular NAMPT levels in body tissues. A possible explanation is the balancing of NAD^+^ levels by *de novo* synthesis through the tryptophan kynurenine pathway. Defects caused by NAD^+^ deficiency can also be reversed by the supply of NMN in the experimental animals [12].

In animal models of AD, improvement of memory impairment and inhibition of Tau phosphorylation was accomplished by the induction of mitophagy. [13]. Significantly, NMN, a NAD^+^ precursor and the product of NAMPT activity, is one of the most powerful inducers of mitophagy [14], and it can occur by multiple mechanisms, which include NAD^+^ dependent proteins SIRT1, SIRT3, SIRT6, SIRT7, SARM1, CD38, IL-1, and more [15]. Mitophagy stimulation by NMN is considered a therapeutic target for AD; therefore, NAMPT activators should be beneficial for pharmacological purposes. Increased NAD^+^ levels were also observed as a consequence of the calorie restriction (CR), which was accompanied by increased SIRT1 expression and SIRT1-mediated activation of α-secretase [16]. Enzymes participating in different steps of the NAD^+^ biosynthetic pathway, including NAMPT, protect axonal functions in cultured dorsal root ganglion neurons.

Similarly, NAD^+^ and its precursors NAM and NMN also have protective effects on axons [17]. *β*-amyloid plaque load and ATP levels significantly improved both in the cortex and hippocampus by administration of NAD^+^ to the APPswe/PS1 ΔE9 animal model [18]. Protein expressions of NAMPT and SIRT1 in the cortex and hippocampus are decreased in APPswe/PS1 ΔE9 mice, which can be reversed by administration of NAD^+^ [18, 19]. The NAD^+^/NADH ratio is lower in this mouse model compared to control brains, as well [18]. Since NAD^+^ levels and NAD^+^/NADH are decreased in these mice after administration of NAMPT inhibitor FK866 treatment, it can be concluded that NAD^+^ levels are generated through the salvage pathway via NAMPT [20]. Therefore, the activation of brain NAMPT can provide benefits for AD treatment.

Since NAMPT inhibitors are also desired due to their potency to limit the growth of cancer cells [21]; low-toxic NAMPT modulators able to penetrate the blood-brain barrier (BBB) and with desirable pharmacokinetic characteristics should be pharmacologically beneficial. The present work shows an *in-silico* approach for screening such types of NAMPT modulators using compounds from traditional African, Chinese, and Russian medicinal plant libraries. Additionally, FDA-approved and FDA-experimental libraries are involved in the screening as well, to create a shortcut for the repurposing of the known and well-characterized compounds.

## Materials and methods

### Collection of bioactive compounds

Potential NAMPT modulators were collected from libraries of bioactive compounds originating from African medicinal plants [22][23][24]. As a second source, a server for drug repurposing, DrugRep was used, which is based on a compound library of traditional Chinese medicinal plants [25]. The DrugRep receptor-based screen was run using traditional Chinese, FDA experimental drug, and FDA-approved drug libraries. Input included NAMPT co-crystallized with the substrate NAM (PDB (Protein Data Bank [26]): 8DSI); NAM and the activator NP-A1-R (TIE) (PDB: 8DSC); NAM and the activator quercitrin (PDB: 8DSE); and NAM and the activator NAT 2-(2-tert-butylphenoxy)-N-(4-hydroxyphenyl) (PDB: 7ENQ). Two NAMPT structures co-crystallized with inhibitors GNE-618 (PDB: 4o13) and 5-nitro-1H-benzimidazole (2ZM, PDB: 4N9C) were also included to monitor inhibitor-specific protein-ligand interactions. Compounds providing interactions with Gibbs free energy (ΔG) better (more negative) than −10 kcal/mol were collected for further selection. A compound from Russian traditional medicine, ononetin, was also tested since it is mentioned in the literature as a NAMPT modulator, but the evidence is missing [27].

### Virtual compound screening

The pharmacological properties of 191 compounds from African medicinal plants and 40 preselected compounds from DrugRep screening were further assessed by SwissADME [28]. Predicted BBB permeability was prioritized at this stage of the screening, and compounds not fulfilling this criterion were excluded before assessment of other pharmacological properties for further easy applications. Their toxicities were predicted by ProTox-II, a web server for the prediction of the toxicity of chemicals [29]. Selected compounds were re-screened using CB-Dock2 [30] by docking into the structures 8DSI [31], 8DSE [31], 8DSC [31], 7ENQ [32], 4o13 [33] and 4N9C [34] to obtain 2D protein-ligand diagrams for interaction analysis. The molecular docking analyses were repeated twice, and the average value is presented. The schema of the workflow of the virtual NAMPT modulator screening was created using Adobe Illustrator 2025.

### Protein-ligand interaction analysis

The analysis of the protein-ligand interactions was performed using 2D diagrams of the interactions between the compounds and the active and allosteric sites of NAMPT. PDB files of protein-ligand interactions were exported from the molecular docking application CB-Dock2 and visualized by a 2D diagram function of BIOVIA Discovery Studio Visualizer (DSV) v.24.1.0.23298 [35] and compared with known interactions of control compounds (NAMPT activators and inhibitors, negative control compounds).

### Chemical structure similarity network

The network was constructed by importing the list of compounds containing their names and SMILES identifiers as a comma-delimited CSV-UTF8 (*.csv) file. The data were imported as a network from the file to Cytoscape v 3.10.2 [36] and analyzed by the cheminformatics application ChemViz2 v.2.0.3 [37]. The Cheminformatics Tools application was used to construct a chemical structure similarity network and to export the pharmacokinetic parameters of the selected compounds. The results were created using a Prefuse force-directed layout, where the node distance and the number of common links define the structural similarity in a fingerprint-like manner.

### Preparation of NAMPT structures for molecular dynamics study

Molecular dynamics (MD) analysis requires NAMPT crystal structures without gaps, which are not available in the PDB. Therefore, the crystal structure of NAMPT was predicted by the ESMFold app [38] operating on the Neurosnap platform [39]. The protein sequence P43490 from Uniprot database; (https://www.uniprot.org/) [40] was used as an input, and the prediction was run with 6 recycles. To model the missing amino acids in the 4o13 structure, Modeller 10.3 software was also used according to the protocol described previously [41]. One thousand models were generated, and the model with the lowest Discrete Optimized Protein Energy (DOPE) value was selected for validation.

### Molecular dynamics analysis

MD simulations were performed using Gromacs 2025.2 [42] with the interface Visual Dynamics [43] for generating scripts. The general AMBER force field compatible with proteins and most of the organic and pharmaceutical compounds was applied [44]. Electrostatic interactions were treated using the particle mesh Ewald (PME) algorithm with a cut-off of 12 Å. Each system was simulated under periodic boundary conditions in a cubic box, whose dimensions were automatically defined, considering 10 Å from the outermost protein atoms in all Cartesian directions. The simulation box was filled with TIP3P water molecules [45]. Subsequently, a two-step energy minimization procedure was performed by 5000 steps of steepest descent and 5000 steps of conjugate gradient or until the system reached a resistance force lower than 1000 kJ.mol^-1^.nm^-1^. Next, initial atomic velocities were assigned using the Maxwell-Boltzmann distribution corresponding to a temperature of 300 K. All systems were subsequently equilibrated during two successive NVT and NPT equilibration simulations with 500 ps for each. After this period, all the systems were simulated with no restraints at 300 K in the Gibbs ensemble with a 1 atm pressure using isotropic coupling. All chemical bonds containing hydrogen atoms were restricted using the LINCS algorithm [46], and the time step was set to 2 fs, saving the trajectories every 10 ps. Finally, independent MD runs of 100 ns for the protein-ligand complex were simulated, and simulation trajectories were analyzed with the GROMACS package tools [42]. Root-mean-square deviation (RMSD) and root-mean-square fluctuation (RMSF) were calculated separately for each system, fitting their heavy atoms and taking the initial structure of the production dynamics as a reference. Hydrogen bonds (H-bonds) were also calculated between protein and ligand complexes. It was considered a hit when the distance between two polar heavy atoms, with at least one hydrogen atom attached, was less than 3.5 Å. Radius of Gyration (RG) and Solvent-Accessible Surface Areas (SASA) were also calculated. The data of the time-dependent radius of gyration (RG) and solvent-accessible surface area (SASA) were compared for different protein-ligand interactions using the y-axis values of the linear trendline at the simulation time of 50 ns.

## Results

The main goal of the present study is to identify new modulators of NAMPT activity among compounds from traditional African, Chinese, and Russian medicinal plants and from libraries of FDA-approved and FDA-experimental medicines. Activators can potentially improve mitochondrial bioenergetics during normal aging and/or conditions of patients with neurodegenerative diseases (AD, PD, HD, ALS) by increasing the cellular NAD^+^/NADH ratio. NAMPT inhibitors are also potentially valuable for the treatment of cancer patients due to their limiting effect on cancer cell proliferation. At this stage, the primary focus of the screening is on the brain, and therefore, the ability to penetrate the BBB was set at the early stage of the screening. Results could represent a valuable starting point for experimental development.

As a basic source of naturally occurring compounds, the libraries originated from African medicinal plants [22][23][24], which included a pan-African natural product library (p-ANAPL) [22]; a library of bioactive compounds from central African flora [23], and a database of phytochemicals from medicinal plant *Kigelia africana* [24] were used. As a second source of potential NAMPT modulators, a fully automated receptor-based server for drug repurposing, DrugRep, was utilized, which contains libraries of FDA-approved and FDA-experimental compounds in addition to compounds from traditional Chinese medicinal plants [25]. Ononetin, originating from the Russian traditional medicinal plant *Ononis spinosa* [27], was also included among the candidates.

The list of 231 natural compounds collected from the available sources was subjected to a pharmacological screening using SwissADME for the prediction of the pharmacokinetic characteristics, medicinal chemistry friendliness, and drug-likeness of the candidate compounds [28]. As the first selection criterion, the predicted ability to penetrate the BBB was chosen for direct effect in the brain (Fig. 1). Characteristics such as high gastrointestinal adsorption

**Fig. 1.**
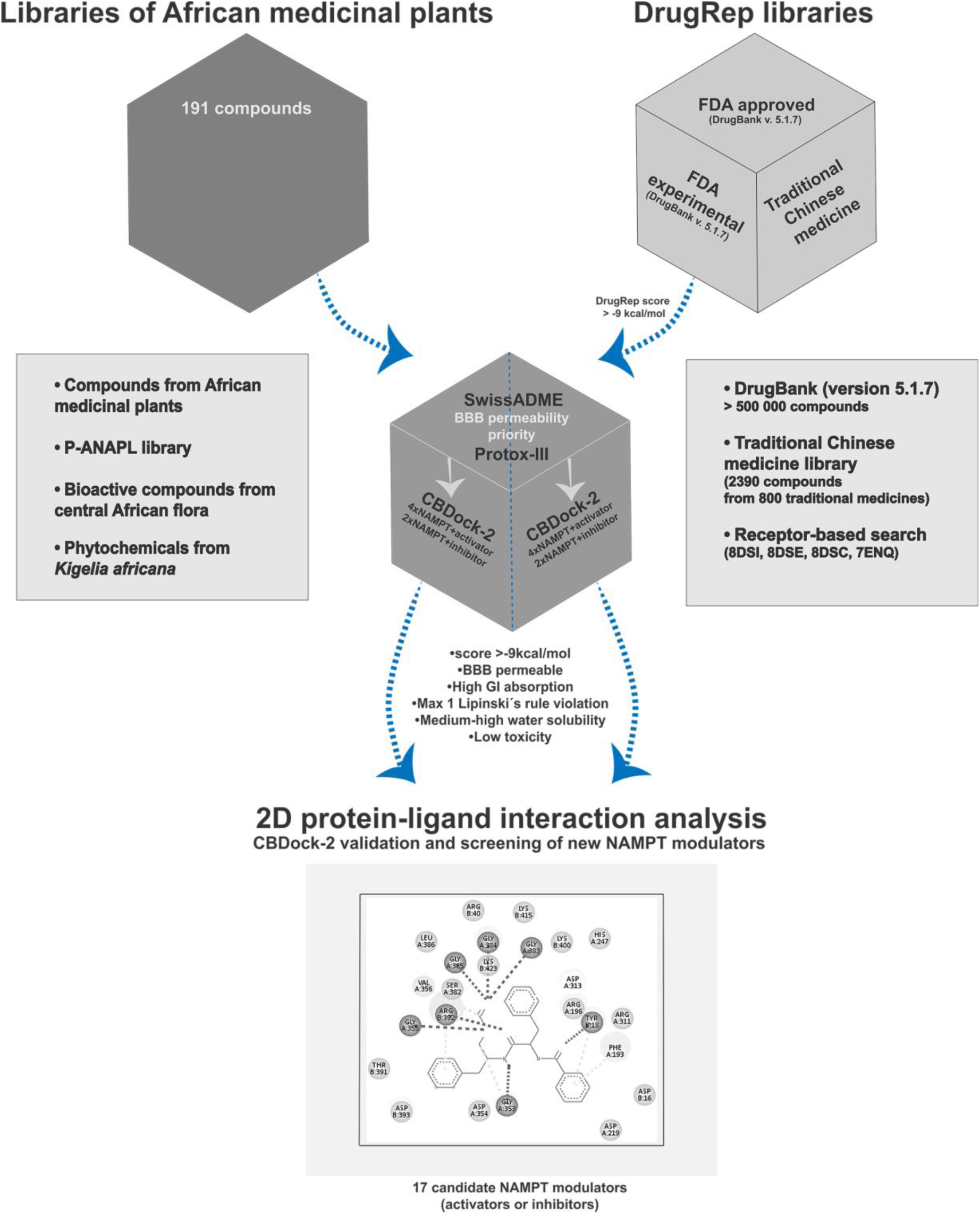
Workflow of the virtual NAMPT modulator screening originated from traditional African, Chinese, and Russian medicinal plants. (1) Firstly, 231 compounds were collected from the libraries of African medicinal plants, DrugRep screening (Chinese traditional medicine), and Russian traditional medicine (ononetin). (2) The compounds were pre-screened by Swiss ADME and Pro-Tox II for BBB permeability, the best predicted pharmacological properties, and low toxicity. Pre-screened compounds were further subjected to molecular docking by CB-Dock2. (3) 2D protein-ligand analysis of the selected and control compounds was performed for validation of the docking method and the comparison of their interactions

(GI), at least moderate water solubility based on SILICOS-IT method [47], and breaking a maximum of one Lipinskís rule, were applied as the next selection criteria of the screening. The compounds fulfilling these basic pharmacological measures were further subjected to the toxicity prediction using Pro-Tox II [29] to exclude highly toxic compounds (Fig. 1).

In general, molecular docking methods suffer from bias towards high molecular weight and lipophilicity of ligands [48]. Here, the highly lipophilic compounds were excluded after the first assessment of BBB permeability, since only water-soluble or moderately water-soluble compounds were selected for further screening. M*_r_* of all new candidates is in the range 318-416 g/mol, which is in agreement with Lipinskís rule, and it decreases the molecular weight effect on the docking scores. The molecular docking was executed by the user-friendly automatic platform CB-Dock2, built on Autodock Vina [30], which yielded 17 compounds with ΔG values lower (more negative) than the threshold of −9 kcal/mol (Fig. 1). The ΔG values of their interactions with NAMPT structures were lower compared to the negative control and comparable with positive control interactions (NAMPT activators and inhibitors), which will be discussed later.

### Validation of the molecular docking method

Molecular docking methods are known to have low consistency; therefore, extensive validation is required to increase the reliability. CB-Dock2 developers promise 85 % accuracy of the obtained results [30], which could provide a good starting point for low-cost screening of bioactive compounds. The ambition of the present work is further improvement of reliability through the use of a panel of positive control compounds with well-characterized binding sites. Therefore, the docking poses of new candidate modulators predicted by CB-Dock2 were compared with known NAMPT modulators through 2-D protein-ligand interaction analysis. Further, negative control compounds were also included to justify the selection threshold of the docking scores. A randomly chosen negative control compound containing multiple aromatic rings demonstrated their roles in the binding to the nucleobase pocket of NAMPT in a similar way as the substrate NAM.

Known interactions of the NAMPT active site and the substrate NAM through the Phe-193/Tyr-18 π−π clamp (with an Arg-311 edge interaction), also called the nucleobase binding site, were used as a primary validation criterion (Tab. 1) [31]. Characterized NAMPT modulators (both activators and inhibitors) containing 4-pyridyl residues are known to bind to both nucleobase and additional “doorsill” orthosteric sites. The active doorsill site binds the substrate NAM and its analogues via the amino acids (AA-s) Asp-219, Ser-241, Val-242, and Ser-275 (Tab. 1) [27]. The Asp-219 is important for the binding of NAM, and His-247 is crucial for autophosphorylation by ATP [49]. Positive allosteric NAMPT modulators, N-PAMs, do not, however, interact with the mentioned nucleobase pocket, but they are bound only to the “rear channel” [31] (Tab. 1) formed by Tyr-188/Gly-185/Ala-379 and Pro-272/Pro-307, with the tolyl group enclosed by His-191/Val-242/Ile-351 and the benzyl group by Arg-349/Val-350. In addition to the interaction with Lys-189, N-PAMs form a loose H-bonding network at the mouth of the rear channel, including Thr-304 (Tab. 1) [27]. These characterized interactions of NAMPT modulators with the active and allosteric sites were applied for the validation of the CB-Dock2 method by 2D protein-ligand interaction analysis and the classification of NAMPT modulator candidates to several binding classes.

**Tab. 1.** An overview of the key amino acids of the active and allosteric sites of NAMPT involved in the binding of known modulators.

| Interaction sites of NAMPT modulators |  | Amino acids |
| --- | --- | --- |
| Active site | Nucleobase pocket | Tyr-18<br>Phe-193 |
|  | Doorsill | Asp-219<br>Ser-241<br>Val-242<br>His-247<br>Ser-275 |
|  | Rear channel | Gly-185<br>Tyr-188<br>Lys-189<br>His-191<br>Val-242<br>Pro-272<br>Thr-304<br>Pro-307<br>Arg-349<br>Val-350<br>Ile-351<br>Ala-379 |

The validation panel of 21 compounds was created from known NAMPT activators, inhibitors, and negative control compounds (Tab. 2). As positive controls, the substrate NAM, the product NMN, and NAMPT activators NAT5-r [32], P7C3 (1-(3,6-dibromo-carbazol-9-yl)-3-phenylamino-propan-2-ol) [50], quercitrin [31], SBI-797812 [51], TIE (NP-A1-R; (3R)-1-[2-(4-methylphenyl)-2H-pyrazolo[3,4-d] pyrimidin-4-yl]-N-{[4-(methylsulfanyl)phenyl]methyl}piperidine-3-carboxamide) (PDB: 8DSC) [31], US11452717 (1-(4-((4-chlorophenyl)sulfonyl)phenyl)-3-(oxazol-5-ylmethyl)urea) [52] and ZN-2-43 ((3S)-N-[(1-benzothiophen-5-yl)methyl]-1-{2-[4-(trifluoromethyl)phenyl]-2H-pyrazolo[3,4-d]pyrimidin-4-yl}piperidine-3-carboxamide) (PDB: 8TPJ) were used [31]. Further, also NAMPT inhibitors 1QR (N-[4-(piperidin-1-ylsulfonyl)benzyl]-1H-pyrrolo[3,2-c]pyridine-2-carboxamide) (PDB: 4KFN) [53], 20M (N-(4-{[4-(pyrrolidin-1-yl)piperidin-1-yl]sulfonyl}benzyl)-2H-pyrido[4,3-e][1,2,4]thiadiazin-3-amine 1,1-dioxide) (PDB: 4LVA; [54]), C5V ((1S,2S)-N-{4-[(1,3-dioxo-1,3-dihydro-2H-isoindol-2-yl)methyl]phenyl}-2-(pyridin-3-yl)cyclopropane-1-carboxamide) (PDB: 6AZJ), FK866 [55], GMX1778 (2-[6-(4-chlorophenoxy)hexyl]-1-cyano-3-pyridin-4-ylguanidine) [56], GNE-617 and GNE-618 [57] [58] were also included in the validation list (Tab. 2).

**Tab. 2.**
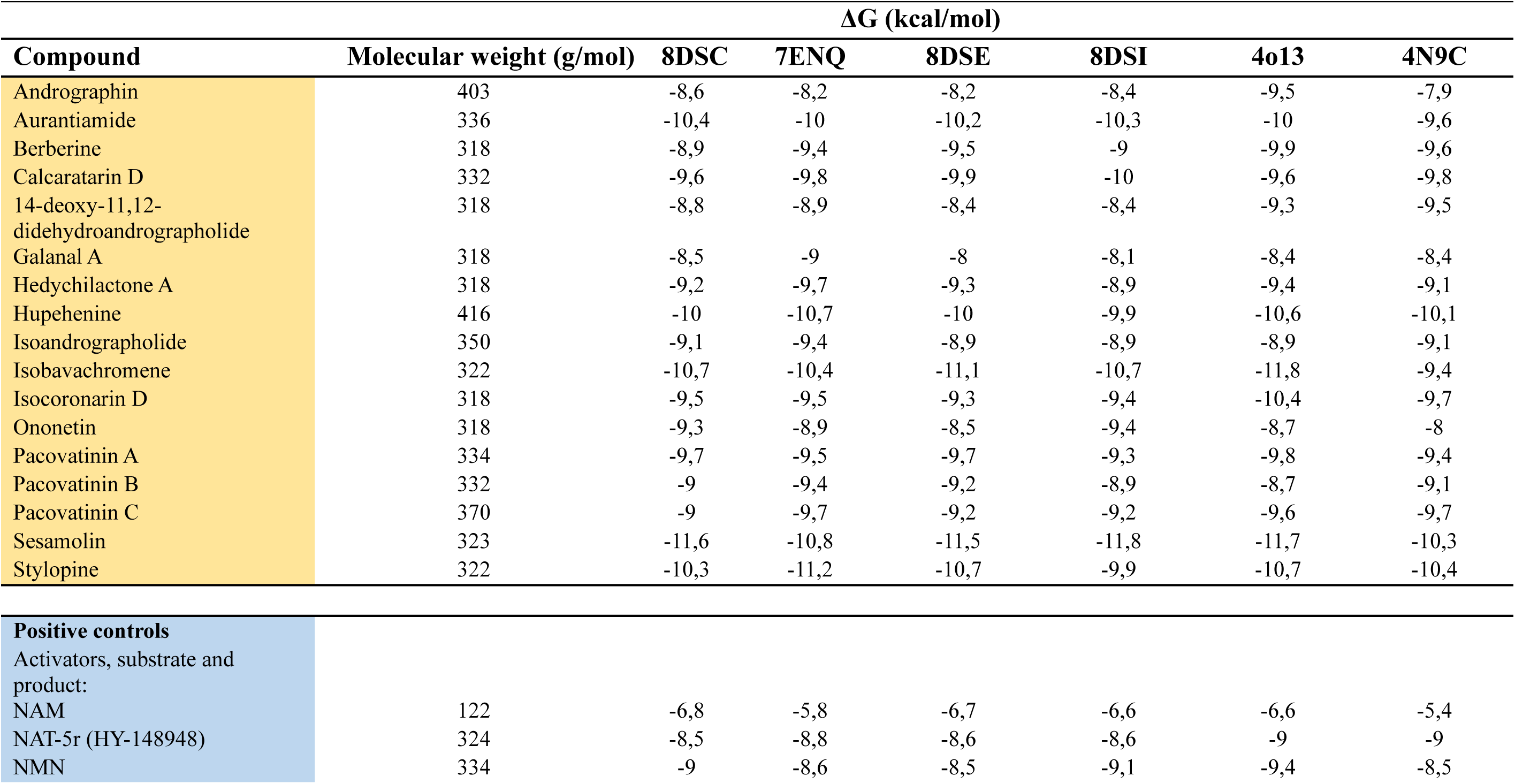
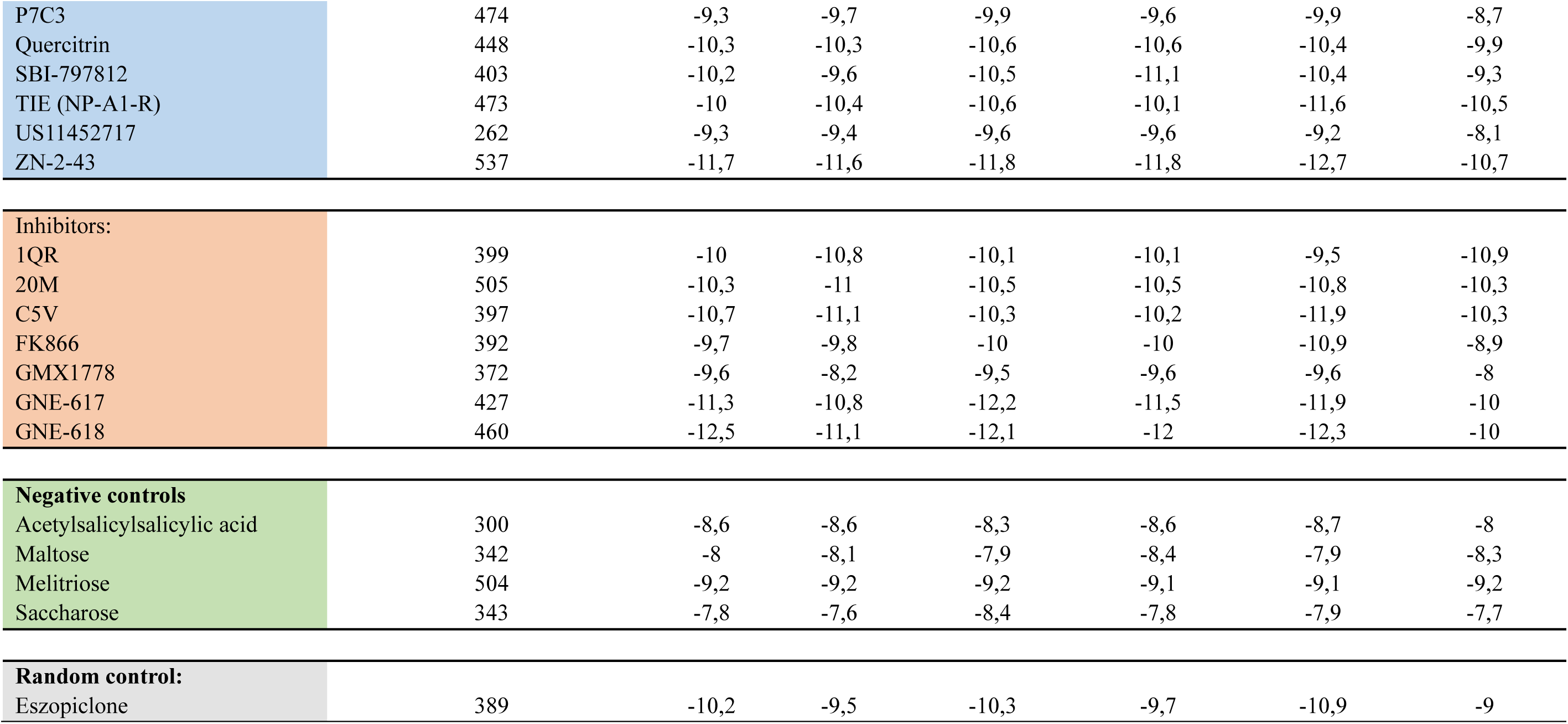
ΔG of the selected candidate NAMPT modulators and control compounds using the tool CB-Dock2. Protein crystal structures: 8DSC (NAMPT+NAM+co-activator NP-A1-R); 7ENQ (NAMPT+activator NAT); 8DSE (NAMPT+NAM+activator quercitrin); 8DSI (NAMPT+NAM); 4o13 (NAMPT+inhibitor GNE-618); 4N9C (NAMPT+inhibitor 5-nitro-1H-benzimidazole)

Firstly, the molecular docking by CB-Dock2 was validated by 2D examination of protein-ligand interactions between the substrate NAM and the active site of NAMPT after NAM re-docking into the structure 8DSI (human NAMPT + NAM). The results displayed all three interactions with the key AA-s (Tyr-18, Phe-193, Asp-219; Tab. 1) (Fig. S1a). A control N-PAM compound TIE (NP-A1-R) was anticipated to bind only to the doorsill and rear channel AA-s (Tab. 1) [31]. Here, the molecular docking results correctly did not show any strong π-π stacked and π-donor hydrogen bonds with essential Phe-193/Tyr-18, nor a hydrogen bond with Asp219, similar to the binding of the substrate NAM (Fig. S1a), which is in agreement with the expected results. According to the CBDock2 prediction, the nucleobase pocket AA-s (Phe-193/Tyr-18), however, interacts with TIE via weaker alkyl bonds. The docking results also showed binding to the doorsill/rear channel Val-242 and the rear channel Tyr-188, Ile-351, and Ala-379, which is consistent with previously reported results. VDW interactions with three additional AA-s involved in the doorsill site of NAMPT were also predicted here (Asp219, Ser241, Ser275) (Tab. 3; Fig. S1b).

**Tab. 3.**
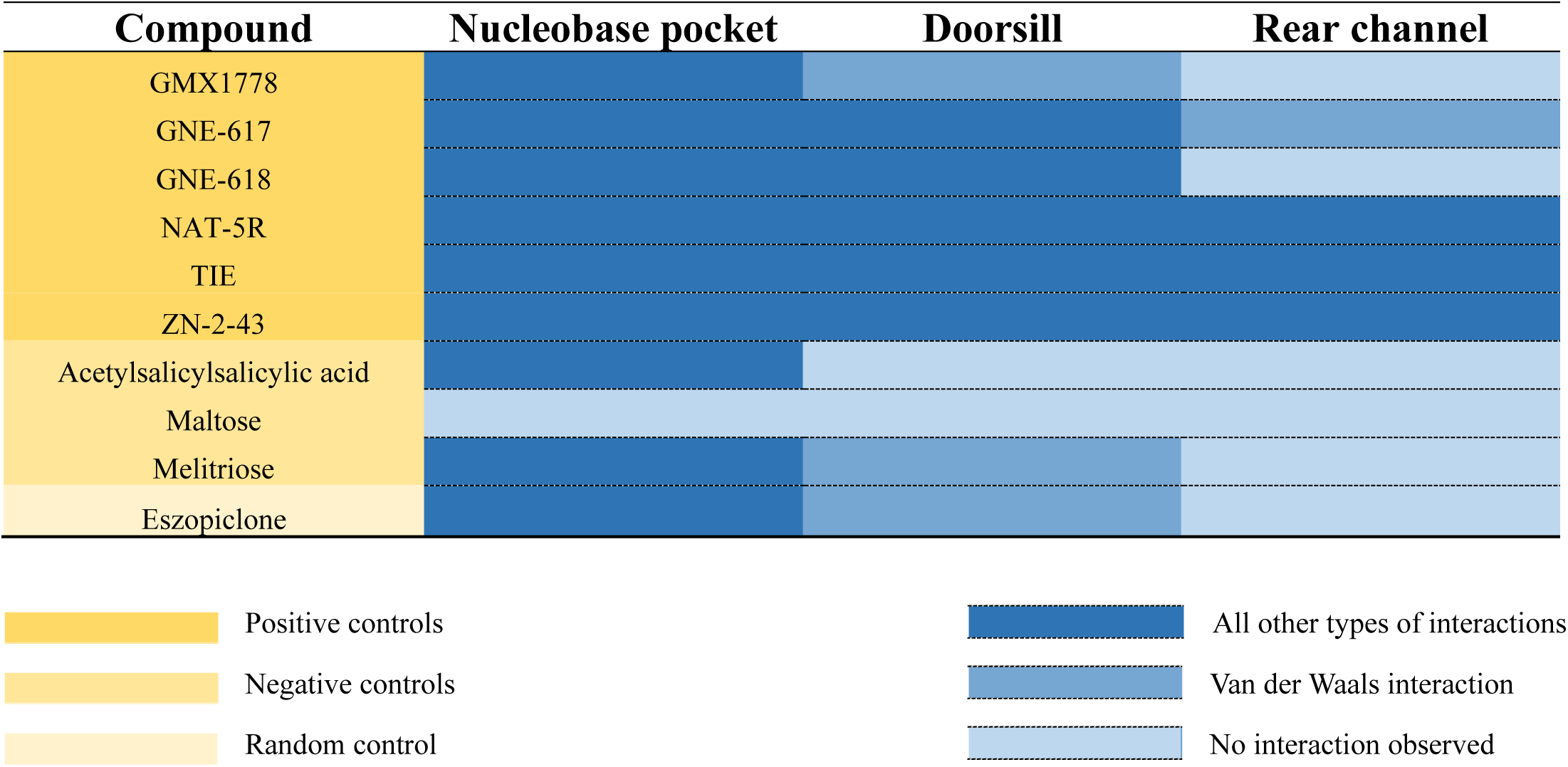
The types of bonds involved in the protein-ligand interactions of positive, negative, and randomly chosen control compounds with the active and allosteric sites of NAMPT.

The common feature of the known strong NAMPT inhibitors GNE-617 and GNE-618 is binding to the Phe-193/Tyr-18 π−π clamp [27]. Here, several known inhibitors and activators interacted with the nucleobase pocket, as it was well documented by docking of GNE-17, GNE-618, and ZN-2-43 (Fig. S1c-e). The CB-Dock2 confirmed π-π stacked interactions of these ligands with the nucleobase pocket, and it generally exhibited highly negative ΔG values reflecting high affinity of their binding to NAMPT. The results of the molecular docking of GNE-617 into 8DSI (ΔG −11,5 kcal/mol) showed strong π-π stacked and π-cation interactions with the nucleobase Phe-193 and Tyr-18, accompanied by π-cation and hydrogen interactions with the doorsill Asp-219 (Tab. 2; Tab. 3; Fig. S1c). GNE-618, when docked into 8DSC (ΔG-12,5 kcal/mol), exhibited π-π stacked interactions with Tyr-18/Phe-193, and hydrogen bonds with the nucleobase Phe-193 and the doorsill Asp-219 (Tab. 3; Fig. S1d). Interactions of another control compound, NAMPT activator ZN-2-43, also exhibited highly negative ΔG in the range from −10,7 to −12,7 kcal/mol with all docked crystal structures. Its interactions with the nucleobase pocket of NAMPT were previously reported to be similar to those of NAM [31]. Here, the interaction with 4o13 (ΔG −12,7 kcal/mol) confirmed π-π stacked interaction with Phe-193 and π-donor hydrogen bond with Tyr-18. Additional interactions were also visualized with AA-s of the doorsill and the rear channel (Tab. 2; Tab. 3; Fig. S1e). ΔG of interactions of another potent activator, NAT-5r (HY-148948) [21] did not reach the values of the best candidate modulators (Tab. 2). However, when docked into the structure 7ENQ ((NAMPT+NAT); ΔG −8,8 Kcal/mol), strong bonds with both the active and allosteric sites were predicted. Bonds with the nucleobase pocket were observed through π-π T-shaped and VDW interactions (Phe-193, Tyr-18). Further, π-alkyl (Val-242), hydrogen (Asp-219), and VDW interactions (Ser-241, Ser-275) of the doorsill and π-alkyl (Arg-349) and VDW interactions (Tyr-188, Lys-189, His-191, Ala-379) of the rear channel AA-s were predicted (Tab. 3, Fig. S1f). Another potent NAMPT inhibitor, GMX1778, docked into the structure 8DSI exhibited strong interactions with Phe-193 by π-π stacked and alkyl bonds and with Tyr-18 by π-donor hydrogen bond (Tab. 3, Fig. S1g), which are identical to nucleobase pocket interactions of the substrate NAM.

Negative controls (acetylsalicylsalicylic acid, maltose, melitriose, saccharose) were also used along with known positive control protein-ligand interactions (Tab. 2, Fig. S2). They were chosen to match the new potential modulators of NAMPT by molecular weight and water solubility. The ΔG values of their interactions were higher than the chosen threshold, except for the interactions of melitriose. The docking of the random control compound acetylsalicylsalicylic acid into 4o13 (ΔG −8,7 kcal/mol) (Tab. 2, Fig. S2a) exhibited the identical types of interactions as the substrate NAM (Tab. 3; Fig. S1a), which seems to occur due to the presence of the aromatic rings. The ΔG values, however, did not reach the values used as the threshold. The observed bonds were also less specific than those of the positive control compound NAT-5r. The docking of maltose as a control compound without aromatic rings did not show any interactions with the key AA-s of the active and or allosteric sites. CB-Dock2 predicted bonds with Arg-392 and Ser-398, which are similar to the binding of the NAMPT product NMN (data not shown). In general, the molecular docking of melitriose with M_r_ >500 g/mol showed only weak VDW interactions with Phe-193 and His-247. When docked into 4o13 and 4N9C, stronger interactions, including a carbon-hydrogen bond with Phe-193 (data not shown) and the hydrogen bond with Tyr-18 (4N9C), were detected (Fig. S2c). Here, the absence of strong types of interactions with the key AA-s of the nucleobase pocket and lack of interactions with the allosteric site indicate low specificity. The ΔG values of melitriose interactions were around the chosen threshold values (Tab. 2), which could also be due to the earlier-mentioned bias by higher M*_r_* compared to other tested compounds.

Further, eszopiclone was randomly chosen as a water-soluble control compound with a molecular weight in the studied range. Its chemical structure contains pyridine, pyrazine, piperidine, and pyrrole rings; therefore, a strong interaction with the nucleobase pocket of NAMPT could be expected. Indeed, the aromatic benzene and pyridine rings exhibited interactions through π-π stacked, π-donor hydrogen, and two π-alkyl bonds with Phe193 and Tyr18. However, no interactions were observed with the doorsill and the rear channel AA-s (Tab. 3; Fig. S2d). The ΔG values of its interactions were much lower than the threshold (from −9 to −10,9 kcal/mol), indicating strong bonds with NAMPT structures. The design of similar structures, including pyridine, pyrazine, piperidine, and pyrrole aromatic rings, could be used for the development of specific NAMPT modulators with desired pharmacological properties. Randomly chosen eszopiclone is also a candidate NAMPT modulator compound; it is, however, already known as a hypnotic agent for treating insomnia with many adverse effects and therefore, is not considered.

To summarize, 2D analysis of TIE interactions with the active and allosteric sites of NAMPT was consistent with the experimental results. The activator ZN-2-43 and strong inhibitors GNE-617, GNE-618, and GMX1778 exhibited interactions with the nucleobase pocket AA-s through π-π stacked, π-π T-shaped, and π−donor hydrogen bonds similar or identical to the binding of the substrate NAM. The ΔG of positive control ligand interactions with NAMPT and their binding patterns with crucial AA-s were also in good agreement with the experimental results. Multiple strong interactions of potent activator NAT-5r with the active and allosteric sites of NAMPT showed, however, the ΔG only around the threshold values and could be expected to be more negative. The analysis also concluded that the presence of multiple aromatic rings can increase the interactions with the nucleobase pocket, similar to NAM, which could be used for the design of specific NAMPT modulators. The ligand without aromatic rings was predicted to interact with AA residues similar to the product of NAMPT enzymatic activity, NMN. 2D protein-ligand interaction analysis of the validation panel of 21 control compounds assisted in creating rules for the selection of new NAMPT modulators.

As a basic rule applied for the selection of new NAMPT modulators is the ΔG of their interactions with NAMPT, which must be more negative than the threshold value of −9 kcal/mol. Further, they should be predicted to interact with the nucleobase pocket (Tyr-18 Phe-193) by strong and the best multiple π-interactions, accompanied by non-VDW interactions with the key doorsill and rear channel AA-s.

### New potential modulators of NAMPT

It is considered a disadvantage that natural compounds have broad specificity since their effect is mediated by numerous protein targets. Therefore, the definition of their mode of action might be useful for the tailored modulation of the target activities. Severe side effects, often present when highly specific synthetic modulators are applied, can be efficiently avoided using selected natural compounds with low toxicity. The selected compounds could provide alternatives for diseases with limited treatment options and many clinical failures, such as Alzheimeŕs and Parkinson’s disease. The present virtual screening method yielded 17 compounds with the interaction ΔG more negative than some known NAMPT modulators and fulfilling basic pharmacological characteristics, including predicted BBB permeability (Tab. 2, Tab. S1). They exhibited specific binding based on the rules identified by 2D analysis of the validation control panel. The candidate NAMPT modulators were further organized according to their interactions with the nucleobase pocket

(Phe-193, Tyr-18) into three subgroups (Tab. 4). Andrographin, aurantiamide, isobavachromene, ononetin, and sesamolin exhibited the strongest interactions with the active site of NAMPT (Tab. 4, group I). Within this group, isobavachromene as the only candidate compound was predicted to interact with both the active and allosteric sites by stronger interactions than VDW bonds and it resulted in one of the lowest ΔG values in the range from −9,4 to −11,8 kcal/mol (Tab. 2). Berberine, calcaratarin D, 14-deoxy-11,12-didehydroandrographolide, galanal A, isoandrographolide, pacovatinin A, pacovatinin C, and stylopine showed weak interactions with the nucleobase pocket through VDW interactions (Tab. 4, Group II) and hedychilactone A, hupehenine, isocoronarin D and pacovatinin B exhibited interactions only with the doorsill and the rear channel AA-s of NAMPT (Tab. 4, Group III). An overview of the selected compounds based on their chemical structure and their pharmacological parameters was constructed through the Cytoscape application Chemviz2 (Fig. 2; Tab. S2). Structurally, 4 compounds from the binding group I (andrographin, isobavachromene, ononetin and sesamolin) are related and created a distinct cluster together with 2 compounds (berberine, stylopine) from the binding group II. The rest of the compounds, excluding aurantiamide and hupehenine formed a larger cluster of structurally related compounds (Fig. 2). Cluster classification served for identification of similarities for possible future structural modifications and search of compound libraries. The interactions of 17 selected NAMPT modulator candidate compounds, representing the main result, will be further discussed in depth.

**Fig. 2.**
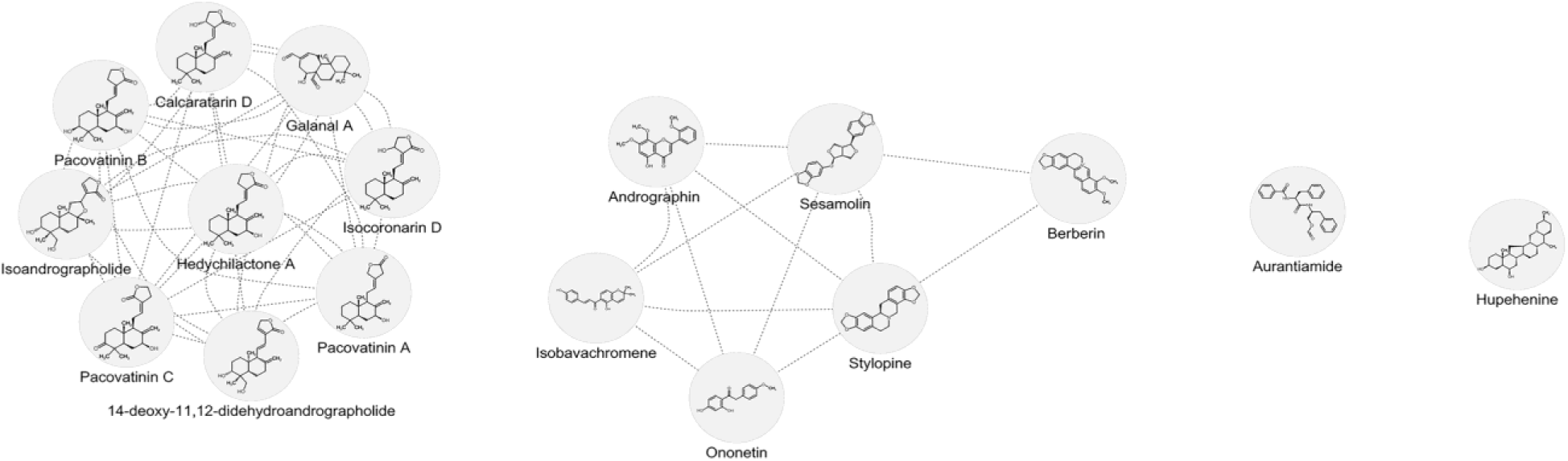
The network of the selected 17 plant NAMPT modulator candidates constructed by importing the list containing chemical names and SMILES identifiers to the cheminformatics application ChemViz2, allowing the construction of the chemical structure similarity network. The results are presented after the Prefuse force-directed layout, where the node distance and the number of common links define the structural similarity in a fingerprint-like manner

**Tab. 4.**
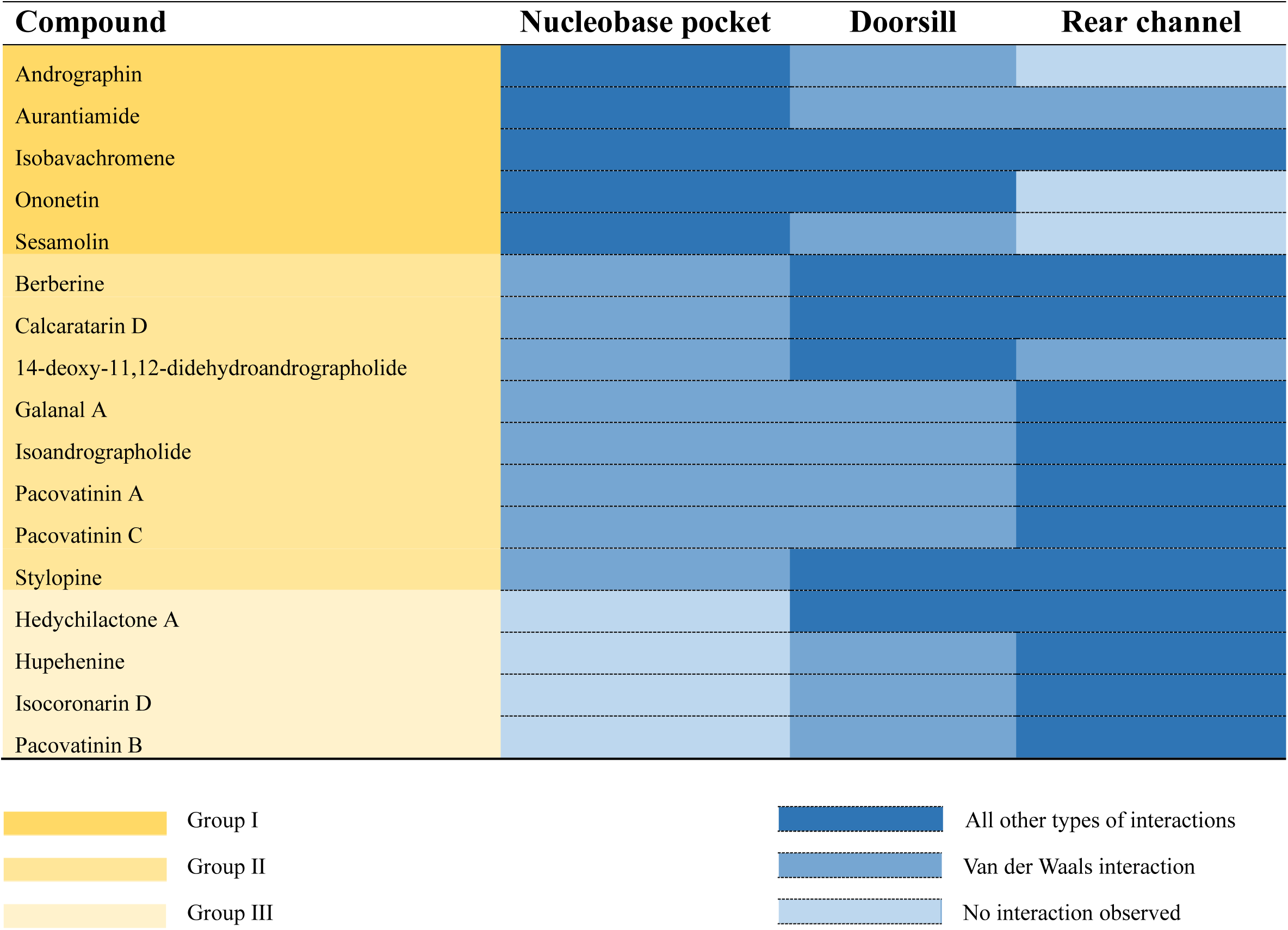
Bonds involved in the protein-ligand interactions of 17 selected candidate NAMPT modulators.

Among new candidate modulators, flavone andrographin originating from *Andrographis paniculata* showed interactions with ΔG lower than the threshold value −9 kcal/mol only when docked into co-crystallized structure 4o13 (NAMPT+GNE-618) (−9,5 kcal/mol) (Tab. 2). Its predicted interaction type is a strong nucleobase pocket binding with the weak doorsill and no rear channel binding (Tab. 4). It is similar to the binding type of NAMPT inhibitor GNE-618, which exhibited, however, stronger interactions with the doorsill AA-s (Tab. 3). In detail, andrographin interactions displayed π-donor hydrogen and π-alkyl bond with Tyr-18; π-π stacked bond with Phe-193, and Van der Waals interactions with the doorsill Asp-219 and His-247 (Fig. 3a; Tab. S3).

**Fig. 3.**
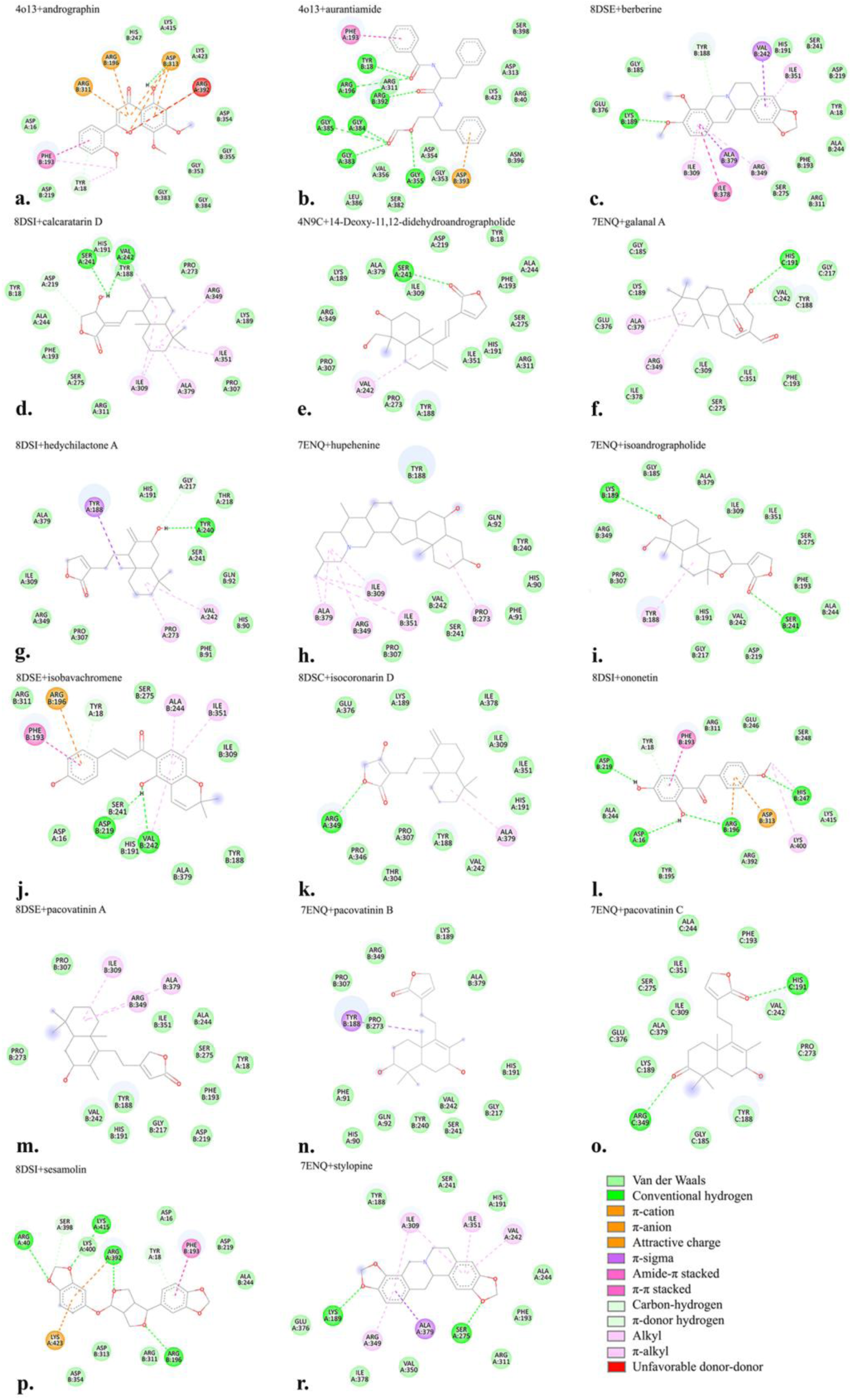
2D diagrams of the protein-ligand interactions between the selected compounds and the key amino acids of the nucleobase pocket, doorsill, and rear channel. The interactions are visualized by BIOVIA Discovery Studio Visualizer after molecular docking of the compounds with the protein crystal structures of NAMPT, employing the molecular docking tool CB-Dock2

Next, the ΔG of the interactions of dipeptide aurantiamide were in the range from −9,6 to −10,4 kcal/mol. Its binding profile with 4o13 was similar to that of andrographin and NAM, with stronger binding to the rear channel AA-s (Fig. 3b; Tab. 4, Tab. S3). Aurantiamide is an exception on the candidate compound list since it is poorly soluble in water (Tab. S1). It can, however, be quickly absorbed in rat tissues after oral administration and eliminated within 4h without accumulation [59], which makes it a good candidate for pharmacological purposes. Another interesting natural compound is benzylisoquinoline berberine, which, when docked into 8DSE showed weak interactions with the nucleobase pocket through VDW bonds. Strong bonds, however, were predicted with the doorsill and rear channel AA-s; therefore, it belongs to the binding group II (Fig. 3c; Tab. 4, Tab. S3). Berberine is known to improve cognitive functions in animal models of AD [60] and its neuroprotective effects provide therapeutic possibilities for AD, PD, and HD treatments [61]. The molecular mechanism of its role in AD could also involve the regulation of amyloid precursor protein (APP) processing to the *β*-amyloid. The oral administration of berberine to AD animal models, rabbits with brain injections of aluminium maltolate, protected the hippocampus from neurodegeneration and decreased BACE1 activity [62]. It functions as a BACE1 inhibitor with IC50 > 100 μM [63], which was used here as an additional control by docking into the structure 7MYI (ΔG −9.7 kcal/mol). Potential neuroprotective effects could also be mediated by the activation of NAMPT activity and the boosting of the NAD^+^/NADH ratio suggested by the current results.

The labdane diterpenoid calcaratarin D exhibited weak interactions with the nucleobase pocket through VDW bonds (Tab. 4). Stronger interactions with the doorsill *via* hydrogen bonds (Ser-241, Val-242), carbon-hydrogen bond (Asp-219), and the rear channel by hydrogen (Tyr-188) and alkyl bonds (Arg-349, Ile-351, Ala-379) were also predicted (Fig. 3d; Tab. 4, Tab. S3). The interactions of another labdane diterpenoid 14-Deoxy-11,12-didehydroandrographolide, included only weak VDW bonds with the nucleobase, doorsill, and rear channel AA-s. The exceptions were the hydrogen (Ser-241) and alkyl (Val-242) interactions at the doorsill (Fig. 3e; Tab. 4; Tab. S3) and the predicted interactions corresponded well to ΔG around the threshold values in the range from −8,4 to −9,5 kcal/mol (Tab. 2). Similarly, the diterpenoid galanal A also exhibited weak VDW interactions with the nucleobase pocket, doorsill, and rear channel accompanied by a few additional alkyl (Arg-349 and Ala-379), carbon-hydrogen (Tyr-188) and hydrogen (His-191) interactions within the rear channel (Fig. 3f; Tab. 4; Tab. S3). The docking poses also corresponded well to ΔG values around the threshold in the range from −8 to −9 kcal/mol (Tab. 2). Another labdane diterpenoid hedychilactone A, also did not exhibit any interaction with the nucleobase pocket of 8DSI (binding group III). It, however, showed interactions with the doorsill through alkyl (Val-242) and VDW (Ser-241) bonds and with the rear channel by π-sigma (Tyr-188) and VDW interactions (His-191, Arg-349, Ala-379) (Fig. 3g; Tab. 4; Tab. S3). The alkaloid hupehenine presented only VDW interactions with the doorsill Ala-379 and alkyl bonds with the rear channel Arg-349, Ile-351, and Ala-379 (Fig. 3h; Tab. 4; Tab. S3). Isoandrographolide, another member of the labdane diterpenoid family, exhibited weak VDW interaction with the nucleobase pocket (Phe-193), single hydrogen interaction with the doorsill (Ser-241), and interactions with the rear channel through the alkyl (Tyr-188) and hydrogen bonds (Lys-189). Additionally, multiple VDW interactions were predicted at both the doorsill and the allosteric site (Fig. 3i; Tab. 4, Tab. S3). These few predicted non-VDW interactions with the crucial AA-s also corresponded well to the ΔG around the threshold values from −8,9 to −9,4 kcal/mol.

Flavone isobavachromene displayed binding to the nucleobase pocket the identical way as NAM, after docking into 8DSE, 8DSI, 7ENQ, and 4o13 (Fig. 3j, Tab. S3). Additional interactions included hydrogen (Asp-219; Val-242) and VDW bonds with the doorsill, alkyl (Ile-351) and multiple VDW interactions with the rear channel (Fig. 3j). It is the only compound among the candidate NAMPT modulators predicted to interact with both active and allosteric sites by stronger bonds than VDW (Group I; Tab. 4). This finding correlates well with decreased ΔG values of its interactions with NAMPT in the range from −9,4 to −11,8 kcal/mol (Tab. 2).

The labdane diterpenoid isocoronarin D belongs to the binding group III since no interaction with the nucleobase pocket was predicted. Further, it showed a weak VDW bond with the doorsill (Val-242) and hydrogen (Arg-349), alkyl (Ala-379), and multiple VDW interactions with the rear channel (Fig. 3k; Tab. 4, Tab. S3). These interactions corresponded to the predicted ΔG values from −9,3 to −10,4 kcal/mol (Tab. 2).

Ononetin, a stilbenoid originating from the traditional Russian medicinal plant *Ononis spinosa* (*Fabaceae*), was previously listed as an NAMPT activator in the work of Velma et al. (2024) [27]; however, the experimental pieces of evidence are missing. Since the compound has low toxicity and is predicted to be BBB-permeable, it was added to the list of tested compounds (Tab. 2). The binding type of group I was predicted for ononetin docked into the structure 8DSI, since the interaction with the nucleobase pocket was identical to that of NAM. The interactions with the doorsill Asp-219 and His-247 were observed through hydrogen bonds (Fig. 3l; Tab. 4; Tab. S3). Labdane terpenoids pacovatinin A-C interacted weakly or not at all with AA-s of the nucleobase pocket and the doorsill AA. Only a few additional interactions with the rear channel of the type π-sigma, alkyl, and hydrogen bonds were predicted, which suggests the modulation of NAMPT activity mostly through the allosteric site (Tab. 4). In detail, pacovatinin A docked into 8DSE displayed weak VDW interactions with AA-s of the active and allosteric sites. Additional alkyl interactions with Arg-349 and Ala-375 located at the rear channel were displayed as well (Fig. 3m; Tab. 4; Tab. S3). Pacovatinin B showed the lowest ΔG −9,4 kcal/mol when docked into 7ENQ (NAMPT+NAT). It exhibited weak VDW with the doorsill (Ser-241, Val-242) and rear channel (Lys-189; His-191; Arg-349, Ala-379), accompanied by only one π-sigma interaction with the rear channel (Tyr-188) AA-s (Fig. 3n; Group II, Tab. 4; Tab. S3). The lowest ΔG of pacovatinin C was also achieved with 7ENQ (NAMPT+NAT), and it corresponded mostly to VDW interactions with the key AA-s and additional hydrogen bonds with the rear channel (His-191 and Arg-349) (Fig. 3o; Tab. 4; Tab. S3). Overall, the interactions of pacovatinin A-C with the protein crystal structures of NAMPT were weak and on the borderline of reliable interactions. Their chemical structures do not contain any aromatic rings interacting strongly with the nucleobase pocket (Tab. S2). The predicted interactions with the doorsill and rear channel AA-s appear, however, more specific compared to the negative controls (Fig. 3m-o; Tab. 4; Tab. S3), and therefore, they are still included in the list of the selected candidate modulators.

Among new NAMPT modulator candidates, the lowest ΔG values in the range from −10,3 to −11,8 kcal/mol were obtained for the interactions of dioxole sesamolin. When docked into the structure 8DSI (NAMPT+NAM), sesamolin displayed the same type of bonds with the nucleobase pocket and the doorsill as the substrate NAM (Fig. 3p; Group I, Tab. 4, and Tab. S3), suggesting a mimicking effect. The observation correlates well with the predicted low ΔG and the presence of two aromatic rings in the chemical structure (Tab. S2). The benzylisoquinoline stylopine also showed a very negative ΔG by docking into all NAMPT co-crystallized structures. When docked into the 7ENQ (ΔG −11,2 kcal/mol), the interaction with the nucleobase pocket was displayed only through VDW interaction with Phe-193. Further, it showed interactions with the doorsill by π-alkyl (Val-242), hydrogen (Ser-275), and VDW (Ser-241) bonds. The rear channel interactions included strong π-alkyl (Arg-349, Ile-351), π-sigma (Ala-379) and VDW (His-191, Tyr-188) bonds (Fig. 3r; Group II; Tab. 4; Tab. S3). The results reflect the effect of the strong π-interactions of stylopine with the doorsill and rear channel of NAMPT, and the resulting binding type group II yielded very low ΔG.

The comparison of binding types between the known and selected candidate modulators did not show any significant differences. As a general observation during the screening of candidate NAMPT modulators, the occurrence of strong π-π stacked, π-sigma, π-alkyl, and π-donor hydrogen bonds with the corresponding AA-s of the active (nucleobase and doorsill) and allosteric site (rear channel) correlated with a more negative ΔG. The incidence of predominantly Van der Waals interactions with only a few additional types of interactions with the crucial AA-s corresponded to ΔG around the threshold value −9 kcal/mol.

Since the natural modulators have multiple targets, they should perform finer tuning of NAMPT activity by their oral administration. The reason to avoid a robust increase in NAMPT activity is the subsequent activation of NAD^+^-dependent enzymes SIRT-s and PARP-s, with boosting of their diverse functions. One of the examples of why this approach must be taken with care is that both NAMPT activity decline and excessive increase have a negative impact on pressure overload-induced heart failure [64]. All members of the NAD^+^ dependent SIRT family perform important tasks in the heart’s functions, and excessive dysregulation of their activities can also have negative effects. An extreme increase in SIRT-s activity is detrimental to cardiomyocytes, which can be prevented by boosting NAMPT-dependent NAD^+^ production by NAM, an inhibitor of SIRT-s [64]. The overexpression of NAMPT also had an undesirable effect in APOE^-/-^ mice, a model of atherosclerosis, where NAMPT promoted atherosclerotic inflammation [65]. These examples, however, illustrate effects due to extreme or even multiple changes of NAMPT expression, and they are the results from models with severe dysregulations. The increase of NAMPT activity in small increments should be a choice, and the natural compounds can provide such alternatives. The development of NAMPT inhibitors for cancer treatment also failed multiple times because of the ability of cancer cells to produce NAD^+^ by alternative pathways [27]. Progress in the development of NAMPT inhibitors should be achieved by the identification of the tumor types based on NAD^+^ production pathways as a part of the precision medicine strategy.

### 3.3 Molecular dynamics simulations

The results of the virtual screening and the final selection of the NAMPT modulators were further supported by the MD simulation analysis. The interactions of three compounds representing the three binding groups I-III (isobavachromene, hedychilactone A and stylopine) were examined along with the positive, negative, and ligand-free controls. Simulations were run using subunit A and B separately to observe the asymmetry of the interactions. The interactions were simulated using the enzymatically active (dimer) and inactive (monomer) NAMPT. Main MD categories were compared to characterize ligand affinity to NAMPT and provide additional evidence of the stability of the protein-ligand complexes.

MD analysis requires NAMPT crystal structures without gaps, which are not available at PDB. All crystal NAMPT structures have a missing short N- and C-terminal peptide, which usually does not influence the analysis; however, missing internal peptides make the MD simulations not feasible. The original structure 4o13 has a missing peptide T44-V52, which was added and used for all MD simulations (assigned in the Fig. S3 as 4o13_gap, further only 4o13). NAMPT structure was also predicted by the ESMFold app, and both structures (Fig. S3A, B) were aligned by Pairwise Structure Alignment (Fig. S3C) [66]. The arrows show the predicted conformation of the missing peptide (Fig. S3A).

Root mean square deviation (RMSD) of the backbone after least square fit into the backbone is a measure of the stability of the protein-ligand complexes. The lowest RMSD values in the range 0,05-0,25 nm confirming the most stable complex with the chain A were observed by binding to isobavachromene (Group I), which were even lower than the positive control compound GNE-618 (Fig. 4a). It corresponded to the average ΔG −11.7 kcal/mol after triple re-docking into the structure 4o13_gap, showing that the adding the missing peptide into the structure 4o13 had only minor effect on the Autodock Vina score (Tab. 2). The RMSD values for binding of stylopine and hedychilactone A were slightly higher than the positive control (4o13+GNE-618), however, lower than the negative control (4o13+maltose). Asymmetry in the results of simulation was obtained for the ligand binding to chain B, with the lowest RMSD values for binding of stylopine and isobavachromene, which were lower than the positive (GNE-618) control (Fig. 4b). ΔG of the triple re-docked stylopine into 4o13_gap structure −10,6 kcal/mol was also nearly identical to ΔG using unmodified 4o13 (Tab. 2). These observations further confirm the stability of the stylopine and isobavachromene complexes with the homodimer NAMPT structure. The RMSD values of hedychilactone A interaction with the chain B were, however, already above the negative control (Fig. 4b). The result also correlates with decreased ΔG of hedychilactone A after docking into 4o13 structure, since the value −8,9 kcal/mol is already higher than the threshold values.

**Fig. 4.**
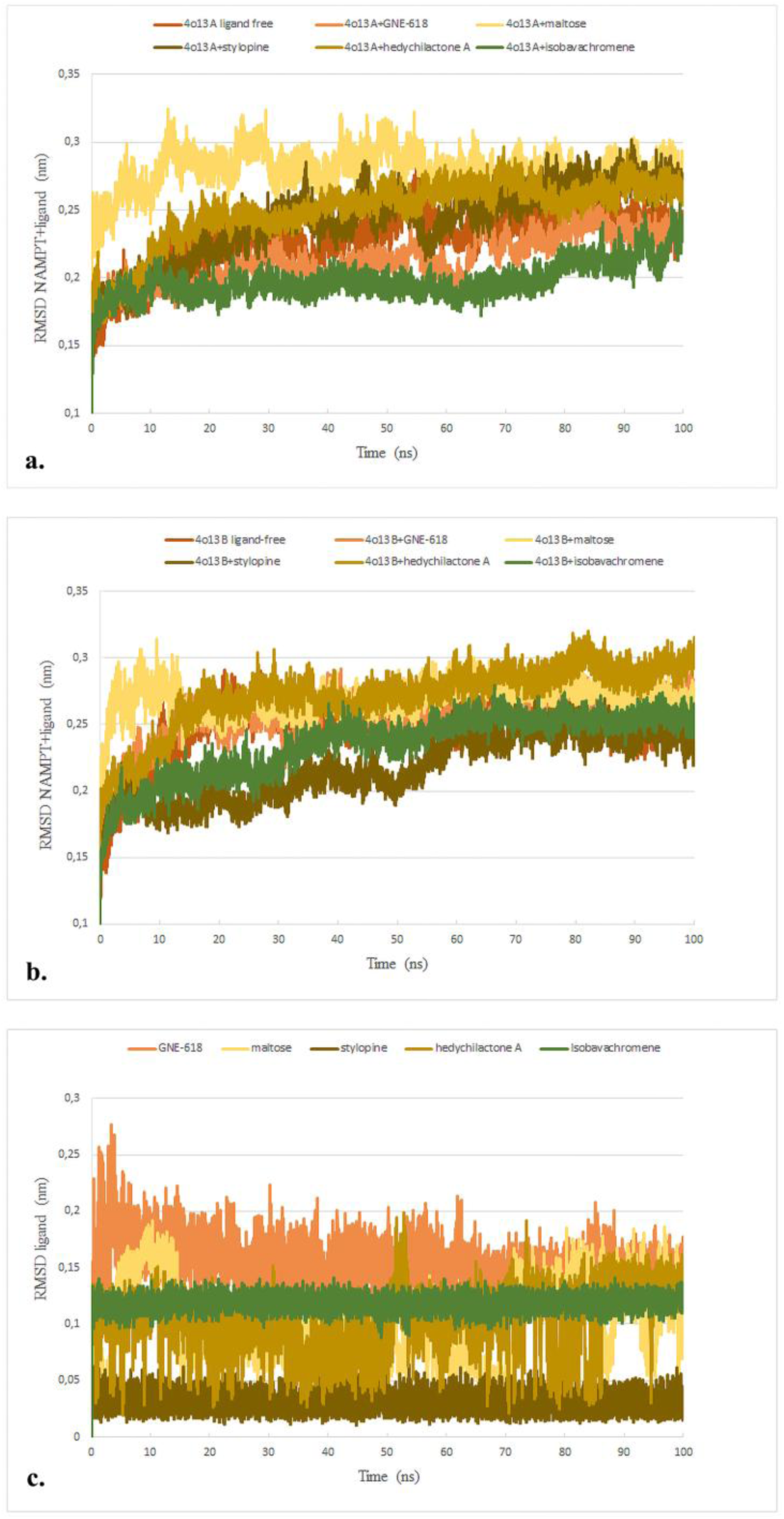
Root mean square deviation (RMSD) of the backbone after least square fit into the backbone of the protein-ligand interactions between three selected potential NAMPT modulators and A. 4o13, chain A; B. 4o13, chain B during a 100 ns simulation period. RMSD of the interactions of ligand-free, positive (4o13+GNE-618), and negative (4o13+maltose) controls were also run in parallel. C. The control simulation runs were performed only with the ligands

Further, RMSD values of the ligand-only simulations varied between ca. 0.02-0,275 nm, and the highest fluctuations and values exhibited negative control maltose, positive control GNE-618, and hedychilactone A, while the values of stylopine and isobavachromene fluctuated to a lesser extent (Fig. 4c). This part of the simulation determines the contribution of the ligand to the values of the 4o13 complexes and might partially explain the failure to confirm the stability of the hedychilactone A and chain B complex. The outcome of this part of the simulation analysis provides further evidence about the specificity of the interactions of new NAMPT modulators isobavachromene, and stylopine. The results also show the asymmetry and reciprocal shift of values for the binding of ligands to both chains. The stability of their complexes with at least one of the 4o13 chains is predicted as comparable or better than the positive control, the strong NAMPT inhibitor GNE-618. Lowest stability was simulated for the hedychilactone A complex with both NAMPT chains. Here, RMSD values were higher than the positive control; however, lower than the negative control (4o13+maltose), when bound to chain A. Binding to chain B was even higher than the negative control; therefore, the RMSD results of MD analysis are only partially supportive for this interaction.

Total radius of gyration (RG) showed minor fluctuations between the values 2,64-2,73 nm for both chains, which suggests high complex stability for binding of all three ligands during the tested period (Fig. 5). RG values of tested compounds were lower than the negative control (4o13+maltose) and comparable or even lower than the positive control (4o13+GNE-618) structures, which indicates higher stability and compactness of the complexes. Simulation of binding to the chain A showed the lowest RG values for stylopine and hedychilactone A (Fig. 5A). However, binding to the chain B displayed the lowest RG values for the interactions of isobavachromene and hedychilactone A (Fig. 5B). RG values around axes X, Y, and Z further demonstrate the difference between binding of all three ligands to chains A and B (Fig. S4). Root mean square fluctuations (RMSF) per residue of NAMPT were also assessed as a measure of rigidity or flexibility of different regions of the protein-ligand complexes. In general, they exhibited high values around the N- and C-termini of all complexes. The results within the residues 51-414 exhibited fluctuations between 0,015-0,43 nm for the binding of all three ligands to chain A or B (Fig. 6A and 6B). The positive control showed the values from 0,04-0,3 nm, and the negative control within the range 0,05-0,3nm; the negative control interactions exhibited, however, a higher frequency of the increased fluctuations. Their extent reflects the presence of flexible regions within the complexes. Within the middle part (AA residues 51-414) of the chain A, isobavachromene interaction showed the lowest values and lowest fluctuations, with only a few higher fluctuations up to 0,3 nm, which suggests the highest stability among the analyzed three complexes. The values were comparable to the positive control (4o13+GNE-618), which exhibited a higher peak at residue 228. The binding of hedychilactone A and stylopine to chain A suffered from one major fluctuation at residue 248, with the values 0,35 and 0,43 nm, respectively, which was higher than the negative control. The negative control showed, however, a higher frequency of high fluctuations when bound to chain A (Fig. 6A). Based on the results of the category RMSF per residue, the only clearly conclusive result provided interaction of isobavachromene with chain A, whose values were the most comparable to the positive control (4o13+GNE-618).

**Fig. 5.**
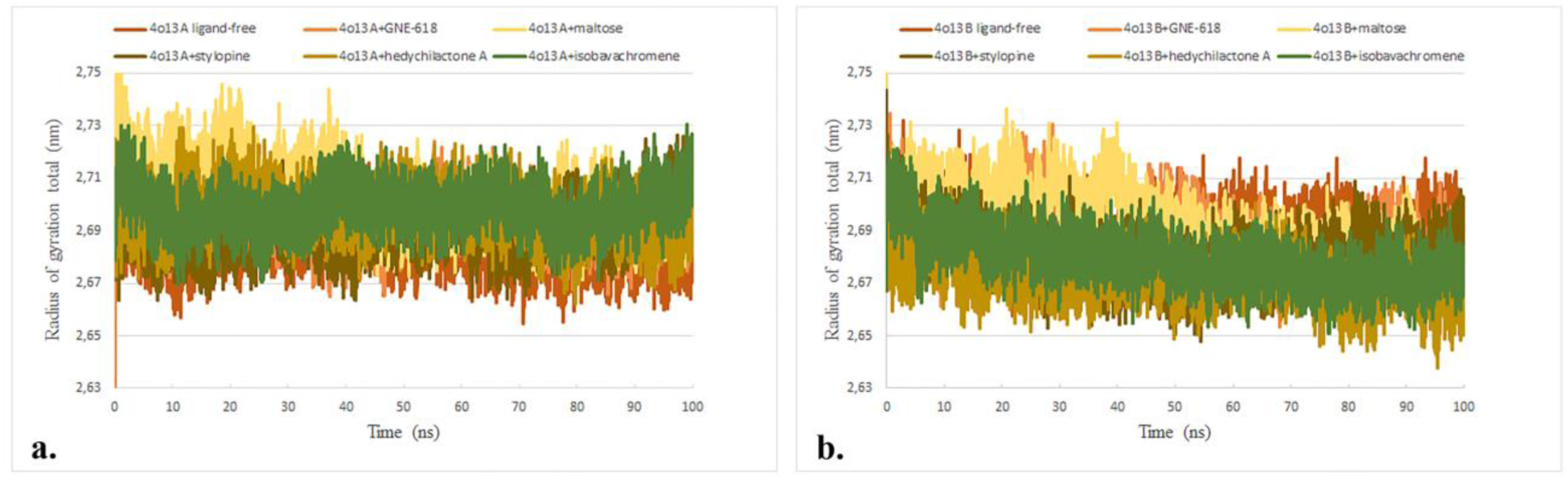
Radius of gyration (RG) of the protein-ligand interactions between three selected potential NAMPT modulators and A. 4o13, chain A; B. 4o13, chain B during a 100 ns simulation period. RG of the interactions of ligand-free, positive (4o13+GNE-618), and negative controls (4o13+maltose) were also simulated in parallel

**Fig. 6.**
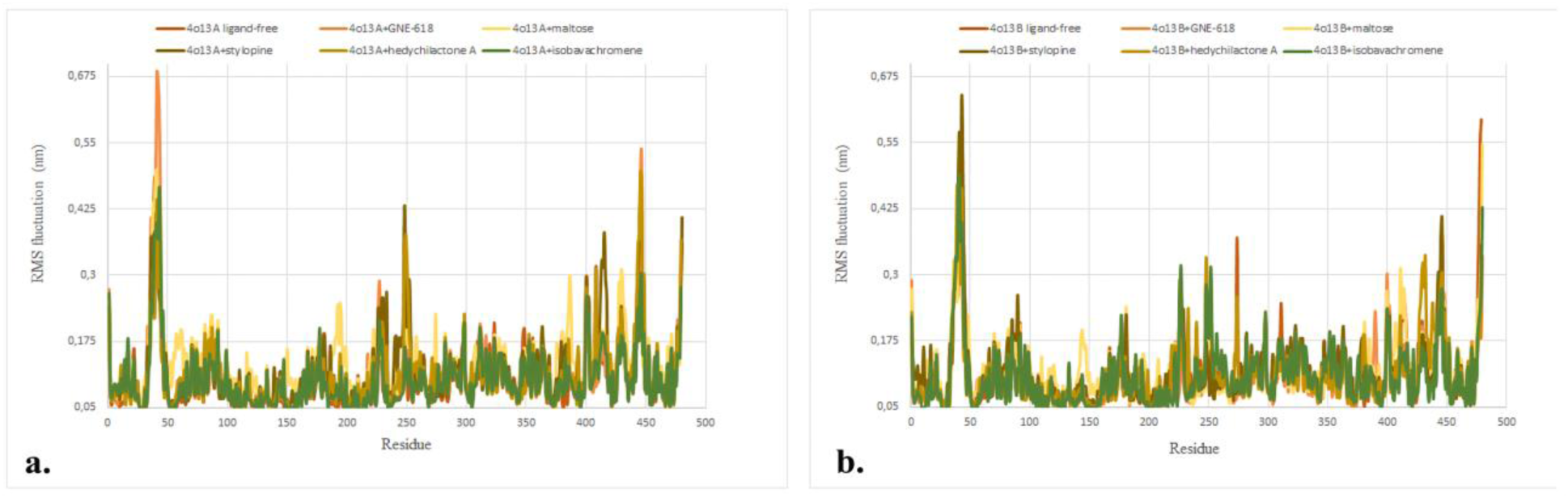
Root mean square (RMS) fluctuations per amino acid (AA) residue of the protein-ligand interactions between three selected potential NAMPT modulators and A. 4o13, chain A; B. 4o13, chain B. RMS fluctuation values per AA residue were also simulated for the interactions of ligand-free, positive (4o13+GNE-618), and negative controls (4o13+maltose)

The comparison of the solvent-accessible surface area (SASA) during the time of the simulation can also reveal information about the stability of the protein-ligand complexes. If the values decrease with the simulation time, it can suggest complex stabilization, allosteric effect, ligand binding, or stabilization of the hydrophobic core due to the ligand effect. This type of profile was the most pronounced by the binding of stylopine and hedychilactone A to chain A and stylopine and isobavachromene to chain B (Fig. 7A, B). The SASA values of the interactions of all three ligands with chain A were lower than the positive and negative control interaction values (Fig. 7A). However, only stylopine and isobavachromene interactions with chain B provided lower SASA values than the negative control interaction (Fig. 7B). The SASA values for hedychilactone A complex within the simulation period showed, however, much higher stabilization than the negative control for binding to chain A, comparable with the positive control. It was not the case for the binding to the chain B, where the values were slightly higher than the negative control (Fig. 7A, B). When the SASA simulation analysis was performed using the homodimer structure 4o13, better stability and compactness of the protein-ligand complexes (data not shown) were obtained compared to the ESMFold NAMPT monomer (Fig. S3). It is in agreement with the enzymatic function of NAMPT as a homodimer, since the active site of the enzyme is located on the interface of two symmetrical units. The nicotinamide moiety interacts through π-stacking interaction with Tyr-18 residue of one and Phe-193 of another chain, thus requiring both NAMPT. chains forming the active site [59]. Therefore, the compounds interacting with the nucleobase pocket (group I) bind to the enzymatically active dimer form, while the compounds belonging to the binding groups II and III could eventually also bind the monomer inactive form.

**Fig. 7.**
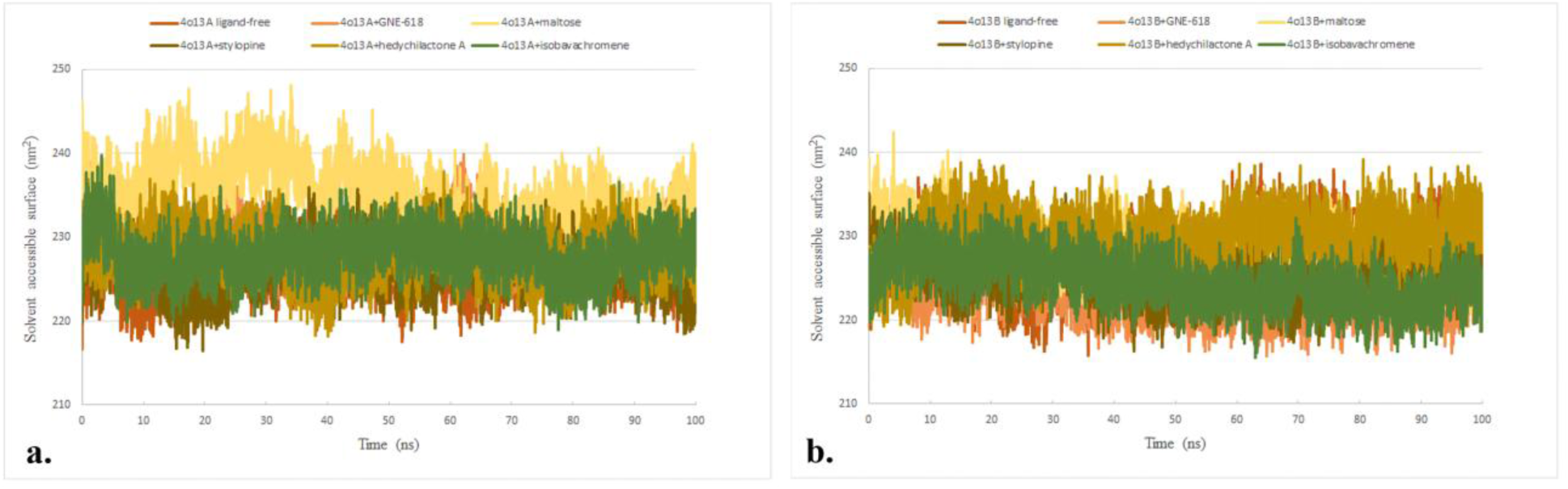
Solvent accessible surface area (SASA) of the protein-ligand interactions between three selected potential NAMPT modulators and A. 4o13, chain A; B. 4o13, chain B during a 100 ns simulation period. SASA of the interactions of ligand-free, positive (4o13+GNE-618), and negative controls (4o13+maltose) were also simulated in parallel

Finally, the total number of the hydrogen bonds (HB) and the number of hydrogen pairs within 0,35 nm were also calculated for the estimation of the binding affinities of tested ligands to NAMPT (Fig. 8). Isobavachromene binding exhibited highest number of the hydrogen bonds (Fig. 8a), which corresponds to the strong interactions with the nucleobase pocket, doorsill and the rear channel (Tab. 4). The number of the hydrogen bonds involved in the interactions fluctuated around 815 during the simulation period. On the other hand, hedychilactone A, belonging to the binding group III, not involving any interactions with the nucleobase pocket (Tab. 4), showed the bottom range of the values around 770 hydrogen bonds in total, which is comparable to the negative control. Interestingly, stylopine (binding group II) provided middle values comparable with the positive control GNE-618, with the fluctuations around 800 hydrogen pairs within the period of the simulation (Fig. 8a). During the tested period, the predicted number of hydrogen pairs formed within 0.35 nm exhibited only minor differences between the simulated protein-ligand interactions (Fig. 8b). All three protein-ligand interactions exhibited values within the range between 4200 pairs (negative control) and 4225 pairs (positive control); therefore, no conclusions are made based on these observations.

**Fig. 8.**
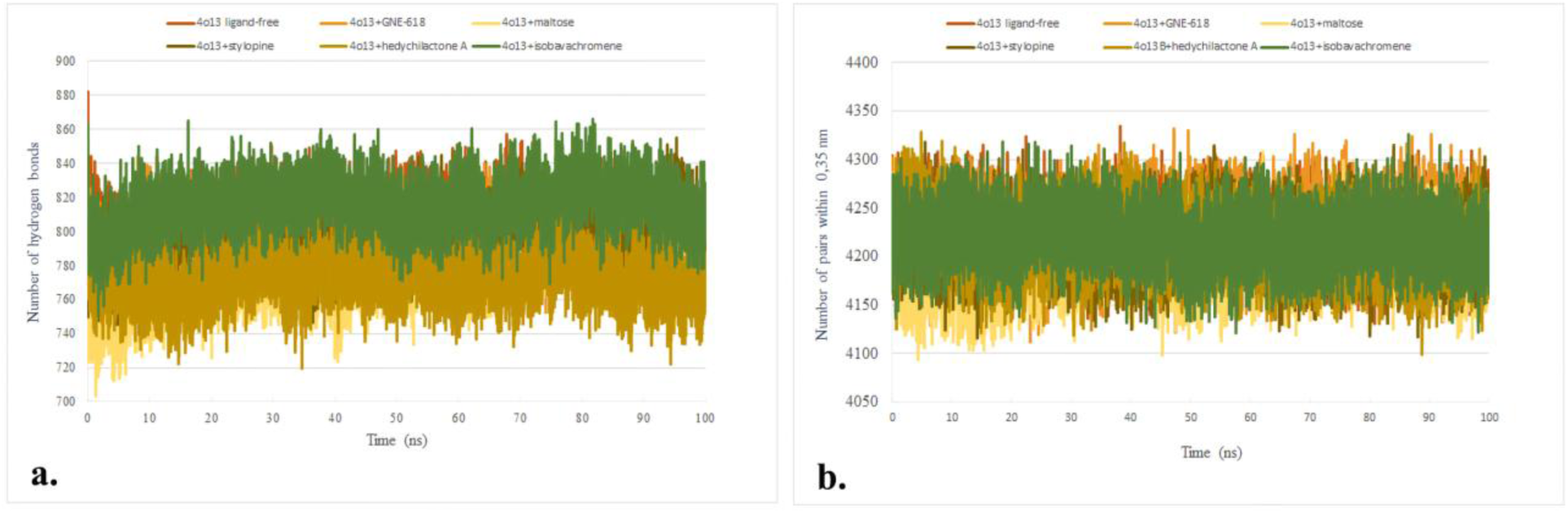
Affinity of three selected potential NAMPT modulators to the dimer 4o13 during a 100 ns simulation period, calculated as A. total number of hydrogen bonds and B. number of hydrogen pairs formed within 0,35 nm. The corresponding values of ligand-free, positive (4o13+GNE-618), and negative controls (4o13+maltose) were also simulated in parallel

The simulation of the total number of hydrogen bonds confirmed the interactions of NAMPT with stylopine and isobavachromene, while the hedychilactone A interaction, since not different from the negative control, was not conclusive based on this category of MD analysis.

## Conclusions

The current study shows a virtual screening method to identify candidate modulators of NAMPT activity, which offers an alternative to experimental screening; however, the final evaluation must always be performed by laboratory experimentation.

NAMPT activators are highly desired for the enhancement of mitochondrial bioenergetics during normal aging and/or for the improvement of conditions of patients with neurodegenerative diseases such as AD, PD, and HD by boosting the cellular NAD^+^/NADH ratio. Increasing NAD^+^ production could also provide aging-related benefits by stimulating the activity of NAD^+^-dependent SIRT-s. Here, however, small stimulatory increments should be chosen because of possible negative effects. Potential NAMPT inhibitors could be used to limit cancer cell growth if the tumor type is dependent on the salvage pathway of NAD^+^ production, which should be determined by the precision medicine approach. Here, the advantage is that the selected compounds do not contribute to additional toxicity to the normal cells compared to most of the approved medications.

Pre-screening of 231 compounds originating from traditional African, Chinese, and Russian medicinal plants, FDA-approved and FDA experimental libraries resulted in the final selection of 17 compounds. These compounds with low toxicity are predicted to pass the BBB, and they fulfil basic pharmacological criteria. They are predicted to interact with the active and/or allosteric sites of NAMPT through specific AA residues similar to characterized NAMPT activators and inhibitors. Based on the type of their binding to the nucleobase pocket of NAMPT, they were divided into three binding groups, and the interactions of one member of each group were confirmed by MD simulation analysis. The interactions of isobavachromene and stylopine from the binding groups I and II, which exhibited more negative ΔG, were confirmed with high confidence using RMDS, RG, RMSF, SASA, and total HB simulation analysis. The interaction of hedychilactone A from the binding group III, which does not form interactions with the nucleobase pocket, and with ΔG slightly below the threshold, was confirmed only by RG and partially by RMSD and SASA simulations. The selected potential modulators of NAMPT activity could serve as a starting point for the development of therapeutic applications by pharmaceutical companies.

## Acknowledgments

The research was funded by Biochemworld Co., Uppsala County, Sweden, which also holds a formal trademark for the published ideas.

## Declarations and statements

### Competing interests

No competing interests to declare.

### Declaration of generative AI and AI-assisted technologies in the writing process

No AI-assisted technologies were used in the present manuscript, neither for figures nor manuscript text generation.

## Abbreviations

AA: amino acid
AD: Alzheimeŕs disease
ALS: amyotrophic lateral sclerosis
BBB: blood-brain barrier
BIOVIA DSV: Biovia Discovery Studio Visualizer
ETC: electron transfer chain
HD: Huntingtońs disease
MD: molecular dynamics
NAM: nicotinamide
NAMPT: nicotinamide phosphoribosyltransferase
NMN: nicotinamide mononucleotide
PD: Parkinsońs disease
PDB: Protein Data Bank
RG: radius of gyration
RMSD: root mean square deviation
RMSF: root mean square fluctuation
SASA: solvent-accessible surface area
VDW: Van der Waals

---

**Fig. S1** 2D diagrams of the protein-ligand interactions between the positive control compounds and the key amino acids of the nucleobase pocket, doorsill, and rear channel. The interactions were visualized by BIOVIA Discovery Studio Visualizer after molecular docking of the compounds with the protein crystal structures of NAMPT using the docking tool CB-Dock2

**Fig. S2** The 2D diagrams of the protein-ligand interactions between the negative and randomly chosen control compounds and the key nucleobase pocket, doorsill, and rear channel amino acids. The interactions were visualized by BIOVIA Discovery Studio Visualizer after molecular docking of the compounds with the protein crystal structures of NAMPT using the docking tool CB-Dock2

**Fig. S3** Preparation of the NAMPT crystal structure for molecular dynamics simulations. A. The NAMPT structure 4o13 was modified by adding the missing peptide T44-V52 (4o13_gap, alternatively 4o13). B. NAMPT structure was also predicted by the ESMFold app. C. Both NAMPT structures were aligned by Pairwise Structure Alignment to visualize modifications. The arrows show the predicted conformation of the missing peptide

**Fig. S4** Radius of gyration (RG) values around axes X, Y, and Z further demonstrate the difference between the binding of three ligands to NAMPT chains A (A., C., E.) and B (B., D., F.) during a 100 ns simulation period. RG values of the interactions of ligand-free, positive (4o13+GNE-618), and negative controls (4o13+maltose) were also simulated in parallel

**Tab. S1** Overview of the selected candidate NAMPT modulators, including their pharmacological characteristics predicted by SwissADME and ProTox-II

**Tab. S2** Pharmacological parameters of 17 selected candidate NAMPT modulators exported from cheminformatics application Chemviz2 run under the Cytoscape 3.10.2 environment

**Tab. S3** The types of bonds involved in the protein-ligand interactions of 17 selected candidate NAMPT modulators with the specific key amino acids of the active and allosteric sites of NAMPT

## Notes

### Competing Interest Statement

The authors have declared no competing interest.

### Summary of Updates

The update presents molecular dynamics data confirming protein-ligand interactions for three compounds across three binding groups defined in the manuscript. The data support and validate previously published findings.

