## Supplementary figures and images for "*In silico* screening of candidate NAMPT modulators for treatment of age-related diseases"

### Fig. S1

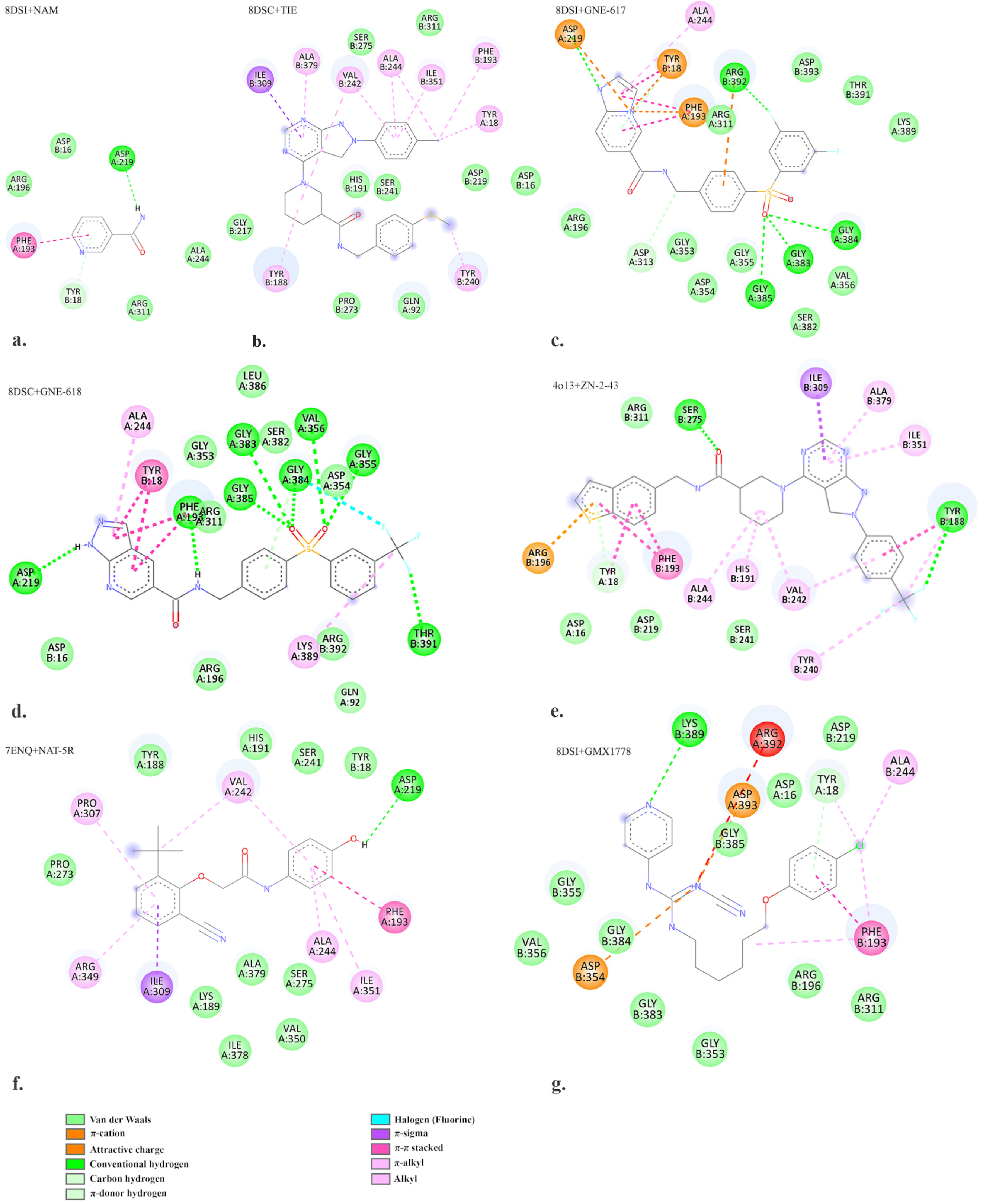

### Fig. S2

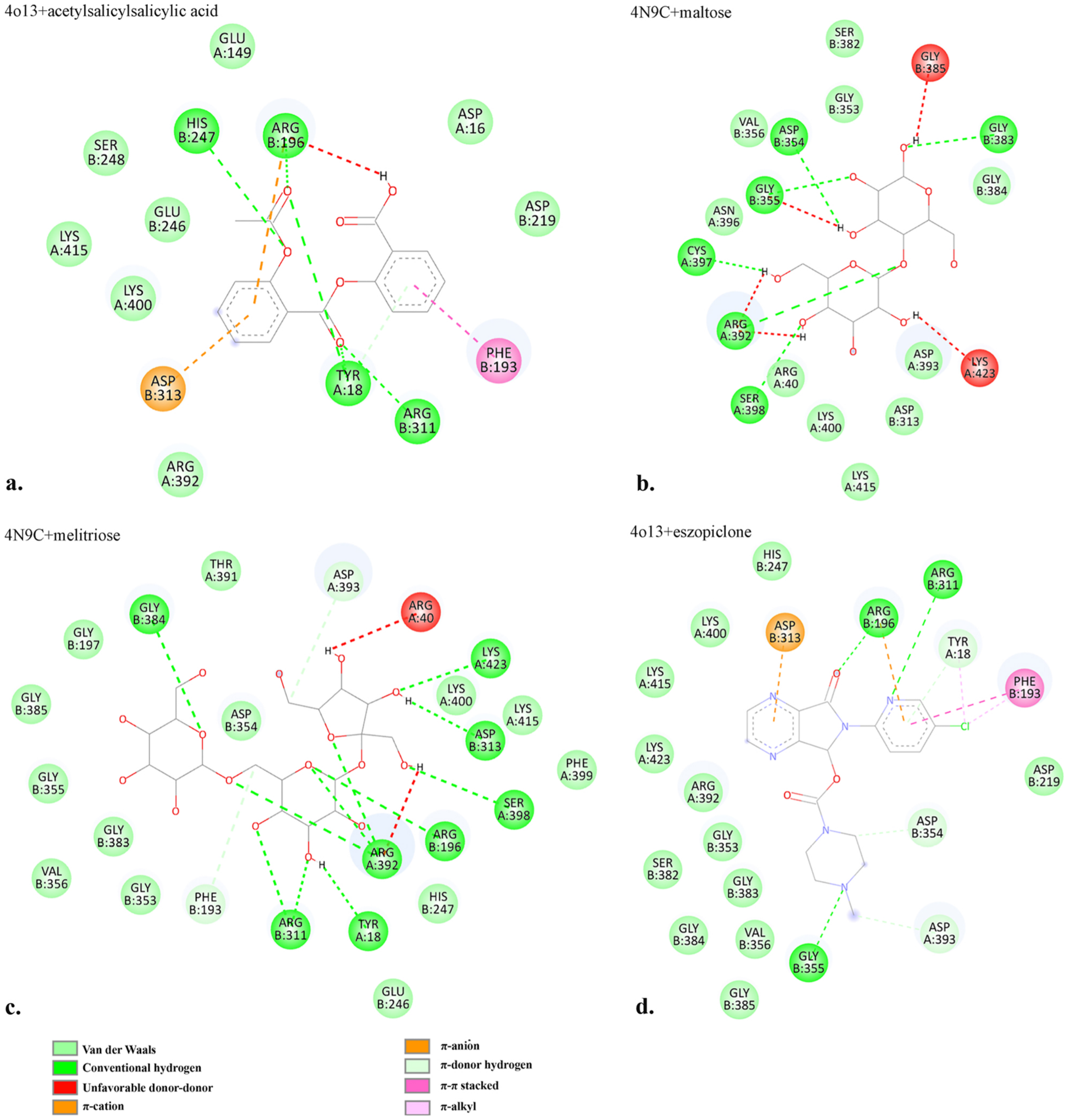

### Fig. S4

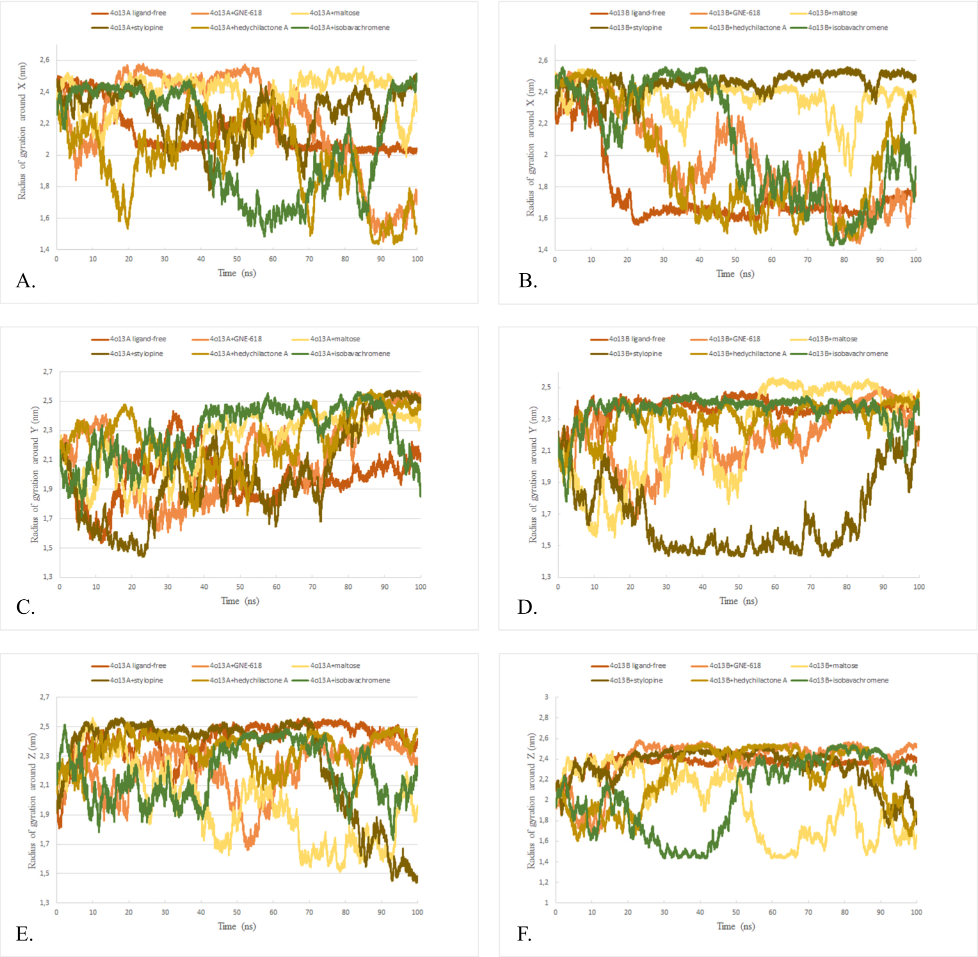
