## Supplementary material for "*In silico* screening of candidate NAMPT modulators for treatment of age-related diseases": Fig. S3

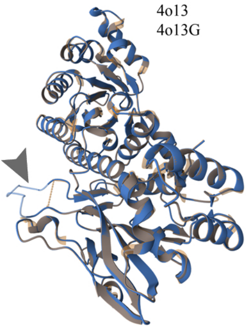


**a.**

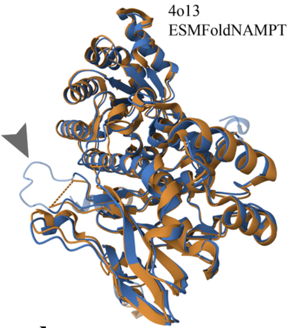


**b.**

```
4O13.A              -------EFNILLATDSYKVTHYKQYPPNTSKVYSYFECREKK---------KYEETVFY    44
NAMPT_ESMFOLD.A     MNPAAEAEFNILLATDSYKVTHYKQYPPNTSKVYSYFECREKKTENSKLRKVKYEETVFY    60
4O13_gap.A          -------EFNILLATDSYKVTHYKQYPPNTSKVYSYFECREKKTENSKLRKVKYEETVFY    53
                           ************************************         ********

4O13.A              GLQYILNKYLKGKVVTKEKIQEAKDVYKEHFQDDVFNEKGWNYILEKYDGHLPIEIKAVP    104
NAMPT_ESMFOLD.A     GLQYILNKYLKGKVVTKEKIQEAKDVYKEHFQDDVFNEKGWNYILEKYDGHLPIEIKAVP    120
4O13_gap.A          GLQYILNKYLKGKVVTKEKIQEAKDVYKEHFQDDVFNEKGWNYILEKYDGHLPIEIKAVP    113
                    ************************************************************

4O13.A              EGFVIPRGNVLFTVENTDPECYWLTNWIETILVQSWYPITVATNSREQKKILAKYLLETS    164
NAMPT_ESMFOLD.A     EGFVIPRGNVLFTVENTDPECYWLTNWIETILVQSWYPITVATNSREQKKILAKYLLETS    180
4O13_gap.A          EGFVIPRGNVLFTVENTDPECYWLTNWIETILVQSWYPITVATNSREQKKILAKYLLETS    173
                    ************************************************************

4O13.A              GNLDGLEYKLHDFGYRGVSSQETAGIGASAHLVNFKGTDTVAGLALIKKYYGTKDPVPGY    224
NAMPT_ESMFOLD.A     GNLDGLEYKLHDFGYRGVSSQETAGIGASAHLVNFKGTDTVAGLALIKKYYGTKDPVPGY    240
4O13_gap.A          GNLDGLEYKLHDFGYRGVSSQETAGIGASAHLVNFKGTDTVAGLALIKKYYGTKDPVPGY    233
                    ************************************************************

4O13.A              SVPAAEHSTITAWGKDHEKDAFEHIVTQFSSVPVSVVSDSYDIYNACEKIWGEDLRHLIV    284
NAMPT_ESMFOLD.A     SVPAAEHSTITAWGKDHEKDAFEHIVTQFSSVPVSVVSDSYDIYNACEKIWGEDLRHLIV    300
4O13_gap.A          SVPAAEHSTITAWGKDHEKDAFEHIVTQFSSVPVSVVSDSYDIYNACEKIWGEDLRHLIV    293
                    ************************************************************

4O13.A              SRSTQAPLIIRPDSGNPLDTVLKVLEILGKKFPVTENSKGYKLLPPYLRVIQGDGVDINT    344
NAMPT_ESMFOLD.A     SRSTQAPLIIRPDSGNPLDTVLKVLEILGKKFPVTENSKGYKLLPPYLRVIQGDGVDINT    360
4O13_gap.A          SRSTQAPLIIRPDSGNPLDTVLKVLEILGKKFPVTENSKGYKLLPPYLRVIQGDGVDINT    353
                    ************************************************************

4O13.A              LQEIVEGMKQKMWSIENIAFGSGGGLLQKLTRDLLNCSFKCSYVVTNGLGINVFKDPVAD    404
NAMPT_ESMFOLD.A     LQEIVEGMKQKMWSIENIAFGSGGGLLQKLTRDLLNCSFKCSYVVTNGLGINVFKDPVAD    420
4O13_gap.A          LQEIVEGMKQKMWSIENIAFGSGGGLLQKLTRDLLNCSFKCSYVVTNGLGINVFKDPVAD    413
                    ************************************************************

4O13.A              PNKRSKKGRLSLHRTPAGNFVTLEEGKGDLEEYGQDLLHTVFKNGKVTKSYSFDEIRKNA    464
NAMPT_ESMFOLD.A     PNKRSKKGRLSLHRTPAGNFVTLEEGKGDLEEYGQDLLHTVFKNGKVTKSYSFDEIRKNA    480
4O13_gap.A          PNKRSKKGRLSLHRTPAGNFVTLEEGKGDLEEYGQDLLHTVFKNGKVTKSYSFDEIRKNA    473
                    ************************************************************

4O13.A              QLNIEL-E---   471
NAMPT_ESMFOLD.A     QLNIELEAAHH   491
4O13_gap.A          QLNIELE----   480
                    ******
```

**c.**
