## Supplementary material for "*In silico* screening of candidate NAMPT modulators for treatment of age-related diseases": Tab. S1

| **Chemical name** | **Classification** | **Lipinski´s rules violation** | **BBB permeability** | **Solubility (SILICOS-IT)** | **Pro-Tox II predicted LD50 (mg/kg)** | **GI absorption** |
| --- | --- | --- | --- | --- | --- | --- |
| Andrographin | Flavone | 0 | Yes | moderately soluble | 5000 | High |
| Aurantiamide | Dipeptide | 0 | Yes | poorly soluble | 550 | High |
| Berberine | Benzylisoquinoline | 0 | Yes | moderately soluble | 200 | High |
| Calcaratarin D | Labdane diterpenoid | 0 | Yes | moderately soluble | 1890 | High |
| 14-Deoxy-11,12-didehydroandrographolide | Labdane diterpenoid | 0 | Yes | soluble | 6060 | High |
| Galanal A | Diterpenoid | 0 | Yes | soluble | 1000 | High |
| Hedychilactone A | Labdane diterpenoid | 0 | Yes | moderately soluble | 1890 | High |
| Hupehenine | Alkaloid | 1 | Yes | soluble | 200 | High |
| Isoandrographolide | Labdane diterpenoid | 0 | Yes | soluble | 1130 | High |
| Isobavachromene | Flavone | 0 | Yes | moderately soluble | 3800 | High |
| Isocoronarin D | Labdane diterpenoid | 0 | Yes | moderately soluble | 1890 | High |
| Ononetin | Stilbenoid | 0 | Yes | moderately soluble | 2000 | High |
| Pacovatinin A | Labdane diterpenoid | 0 | Yes | moderately soluble | 2000 | High |
| Pacovatinin B | Labdane diterpenoid | 0 | Yes | soluble | 1890 | High |
| Pacovatinin C | Labdane diterpenoid | 0 | Yes | soluble | 57 | High |
| Sesamolin | Dioxole | 0 | Yes | moderately soluble | 1500 | High |
| Stylopine | Benzylisoquinoline | 0 | Yes | moderately soluble | 940 | High |
