## Supplementary material for "*In silico* screening of candidate NAMPT modulators for treatment of age-related diseases": Tab. S2

| **Compound** | **Wiener polarity** | **Aromatic ring count** | **Zagreb index** | **Topological polar surface area** | **XlogP** | **AlogP** | **H bond acceptors** | **H bond donors** |
| --- | --- | --- | --- | --- | --- | --- | --- | --- |
| Andrographin | 44 | 3 | 126 | 74.22 | 74.22 | 2.968 | 1 | 1 |
| Aurantiamide | 43 | 3 | 154 | 84.50 | 84.50 | 3.531 | 6 | 2 |
| Berberine | 47 | 3 | 144 | 40.80 | 40.80 | 2.003 | 0 | 0 |
| Calcaratarin D | 42 | 0 | 128 | 46.53 | 46.53 | 1.336 | 3 | 1 |
| 14-deoxy-11,12-didehydroandrographolide | 46 | 0 | 132 | 66.76 | 66.76 | 0.717 | 4 | 2 |
| Galanal A | 52 | 0 | 130 | 54.37 | 54.37 | 0.211 | 3 | 1 |
| Hedychilactone | 42 | 0 | 128 | 46.53 | 46.53 | 1.529 | 3 | 1 |
| Hupehenine | 64 | 0 | 182 | 43.70 | 43.70 | -1.54 | 3 | 2 |
| Isoandrographolide | 52 | 0 | 148 | 75.99 | 75.99 | 0.104 | 5 | 2 |
| Isobavachromene | 38 | 2 | 128 | 66.76 | 66.76 | 3.975 | 1 | 2 |
| Isocoronarin D | 42 | 0 | 128 | 46.53 | 46.53 | 1.336 | 3 | 1 |
| Ononetin | 27 | 2 | 94 | 66.76 | 66.76 | 2.845 | 1 | 2 |
| Pacovatinin A | 41 | 0 | 128 | 46.53 | 46.53 | 1.308 | 3 | 1 |
| Pacovatinin B | 46 | 0 | 134 | 66.76 | 66.76 | 0.956 | 4 | 2 |
| Pacovatinin C | 46 | 0 | 134 | 63.60 | 63.60 | 1.525 | 4 | 1 |
| Sesamolin | 42 | 2 | 158 | 64.61 | 64.61 | 2.234 | 2 | 0 |
| Stylopine | 44 | 2 | 146 | 40.16 | 40.16 | 2.877 | 1 | 0 |
