## Supplementary material for "*In silico* screening of candidate NAMPT modulators for treatment of age-related diseases": Tab. S3

| **Compound** | | **π-π stacked** | **π-donor hydrogen** | | **π-sigma** | | **π-alkyl** | | **Alkyl** | | **Carbon-hydrogen** | | **Hydrogen** | | **Van der Waals** |
| --- | --- | --- | --- | --- | --- | --- | --- | --- | --- | --- | --- | --- | --- | --- | --- |
| Andrographin | | Phe-193 | Tyr-18 | |  | |  | | Tyr-18; Phe-193 | |  | |  | | Asp-219; His-247 |
| Aurantiamide | | Phe-193 | Tyr-18 | |  | |  | |  | |  | |  | | Asp-219 |
| Berberine | |  |  | | Val-242 | | Arg-349 Ile-351 | |  | | Tyr-188 | | Lys-189 | | Tyr-18; Phe-193 |
|  | |  |  | | Ala-379 | |  | |  | |  | |  | | Asp-219; Se-241; Ser-275 |
|  | |  |  | |  | |  | |  | |  | |  | | His-191 |
| Calcaratarin D | |  |  | |  | |  | | Val-242 | | Asp-219 | | Ser-241; Val-242 | | Tyr-18; Phe-193 |
|  | |  |  | |  | |  | | Arg-349; Ile-351; Ala-379 | |  | |  | | Ser-275 |
|  | |  |  | |  | |  | |  | |  | |  | | Tyr-188; Lys-189; His-191 |
| 14-deoxy-11,12-didehydroandrographolide | |  |  | |  | |  | | Val-242 | |  | | Ser-241 | | Tyr-18; Phe-193 |
|  | |  |  | |  | |  | |  | |  | |  | | Asp-219; Ser-275 |
|  | |  |  | |  | |  | |  | |  | |  | | Tyr-188; Lys-189; His-191; |
|  | |  |  | |  | |  | |  | |  | |  | | Pro-307; Ile-351; Arg-349; Ala-379 |
| Galanal A | |  |  | |  | |  | | Arg-349 Ala-379 | | Tyr-188 | | His-191 | | Phe-193 |
|  | |  |  | |  | |  | |  | |  | |  | | Val-242; Ser-275 |
|  | |  |  | |  | |  | |  | |  | |  | | Lys-189; Ile-351 |
| Hedychilactone A | |  |  | | Tyr-188 | |  | | Val-242 | |  | |  | | Ser-241 |
|  | |  |  | |  | |  | |  | |  | |  | | His-191; Arg-349; Ala-379 |
| Hupehenine | |  |  | |  | | Arg-349 Ile-351 Ala-379 | |  | |  | |  | | Ser-241; Val-242 |
|  | |  |  | |  | |  | |  | |  | |  | | Tyr-188 |
| Isoandrographolide | |  |  | |  | |  | | Tyr-188 | |  | | Ser-241 | | Phe-193 |
|  | |  |  | |  | |  | |  | |  | | Lys-189 | | Asp-219; Val-242; Ser-275 |
|  | |  |  | |  | |  | |  | |  | |  | | Gly-185; His-191; Arg-349; Ile 351; Ala-379 |
| Isobavachromene | | Phe-193 | Tyr-18 | |  | |  | | Ile-351 | |  | | Asp-219 Val-242 | | Ser-241; Ser-275 |
|  | |  |  | |  | |  | |  | |  | |  | | Tyr-188; His-191; Ala-379; |
| Isocoronarin D | |  |  | |  | |  | | Ala-379 | |  | | Arg-349 | | Val-242 |
|  | |  |  | |  | |  | |  | |  | |  | | Tyr-188; Lys-189; His-191; Pro-307; Ile-351; |
| Ononetin | | Phe-193 | Tyr-18 | |  | |  | |  | |  | | Asp-219 His-247 | |  |
| Pacovatinin A | |  |  | |  | |  | | Arg-349 Ala-379 | |  | |  | | Tyr-18; Phe-193 |
|  | |  |  | |  | |  | |  | |  | |  | | Asp-219; Val-242; Ser-275; |
|  | |  |  | |  | |  | |  | |  | |  | | Tyr-188; His-191; Ile-351 |
| Pacovatinin B | |  |  | | Tyr-188 | |  | |  | |  | |  | | Ser-241; Val-242; |
|  | |  |  | |  | |  | |  | |  | |  | | Lys-189; His-191; Arg-349; Ala-379 |
| Pacovatinin C | |  |  | |  | |  | |  | |  | | His-191; Arg-349 | | Phe-193 |
|  | |  |  | |  | |  | |  | |  | |  | | Val-242; Ser-275 |
|  | |  |  | |  | |  | |  | |  | |  | | Tyr-188; Lys-189; Ile-351; Ala-379 |
| Sesamolin | | Phe-193 | Tyr-18 | |  | |  | |  | |  | |  | | Asp-219 |
| Stylopine | |  |  | | Ala-379 | |  | | Val-242 | |  | | Ser-275 | | Phe-193 |
|  | |  |  | |  | |  | | Arg-349; Ile-351 | |  | | Lys-189 | | Ser-241 |
|  | |  |  | |  | |  | |  | |  | |  | | Tyr-188; His-191 |
|  | Nucleobase pocket | | |  | |  | |  | |  | |  | |  | |
|  | Doorsill | |  | |  | |  | |  | |  | |  | |  |
|  | Rear channel | |  | |  | |  | |  | |  | |  | |  |
